# *Clostridium innocuum*, an opportunistic gut pathogen, inactivates host gut progesterone and arrests ovarian follicular development

**DOI:** 10.1101/2024.03.15.585140

**Authors:** Mei-Jou Chen, Chia-Hung Chou, Tsun-Hsien Hsiao, Tien-Yu Wu, Chi-Ying Li, Yi-Lung Chen, Kuang-Han Chao, Tzong-Huei Lee, Ronnie G Gicana, Chao-Jen Shih, Guo-Jie Brandon-Mong, Yi-Li Lai, Po-Ting Li, Yu-Lin Tseng, Po-Hsiang Wang, Yin-Ru Chiang

## Abstract

**Highlights:**

- We identified *Clostridium innocuum* as a key player in gut progesterone metabolism.
- Progesterone is converted into epipregnanolone with negligible progestogenic activity.
- We identified the enzyme and mechanisms of microbial epipregnanolone production.
- *C. innocuum* caused decreased serum progesterone and follicular arrest in female mice.
- *C. innocuum* is a causal factor of progesterone resistance in women taking progesterone.

Graphical Abstract

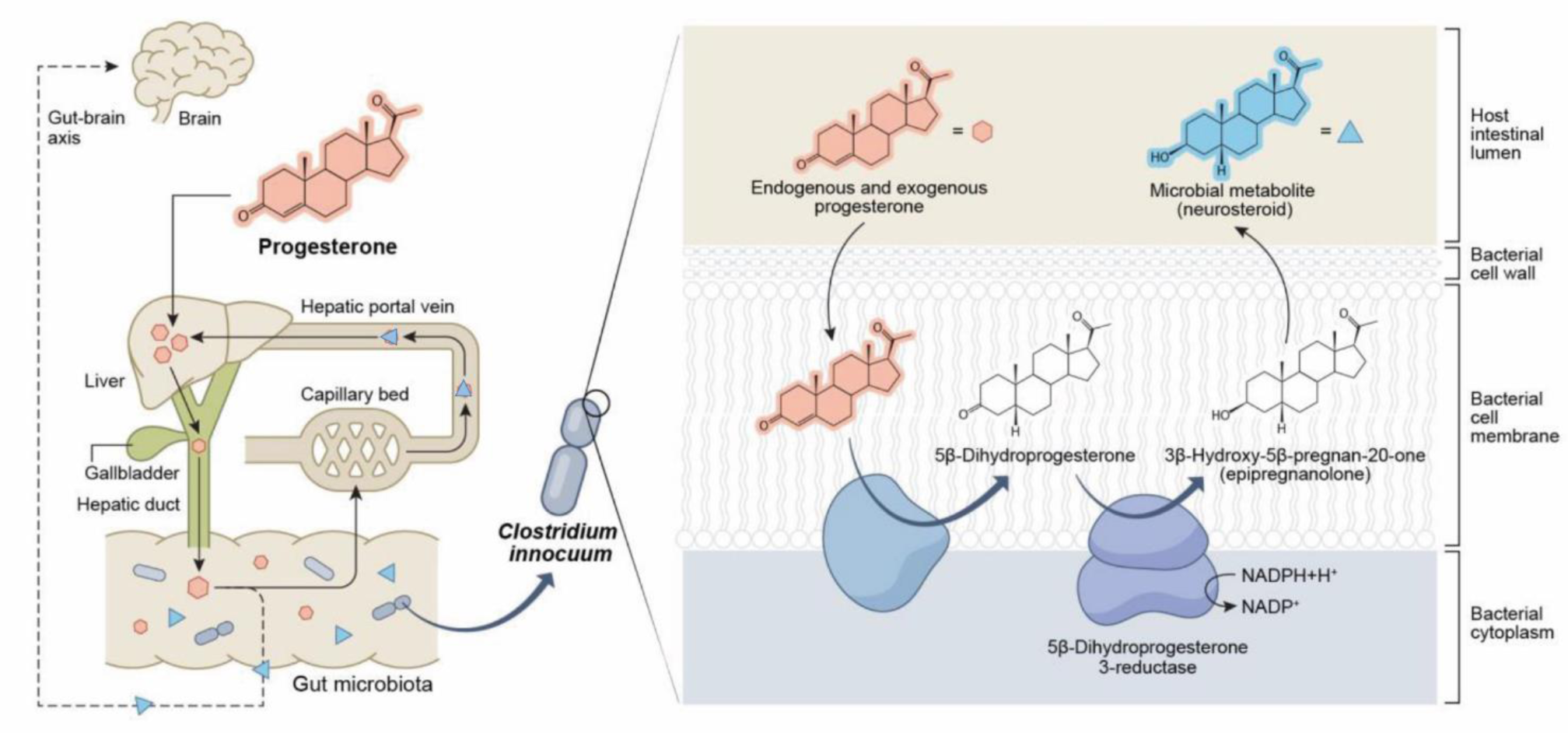

**In brief:** Chen et al. identified *Clostridium innocuum* as a major species involved in gut progesterone metabolism, with epipregnanolone as the main product, and elucidated the molecular mechanisms. *C. innocuum* inactivates gut progesterone in female mice, leading to decreased circulating progesterone levels. *C. innocuum* is also a causal factor of follicular arrest.

Levels of progesterone, an endogenous female hormone, increase after ovulation; progesterone is crucial in the luteal phase to maintain successful pregnancy and prevent early miscarriage. Both endogenous and exogenous progesterone are recycled between the liver and gut; thus, the gut microbiota regulate host progesterone levels by inhibiting enterohepatic progesterone circulation. Our data indicated *Clostridium innocuum* as a major species involved in gut progesterone metabolism in women with infertility. *C. innocuum* converts progesterone into the neurosteroid epipregnanolone (with negligible progestogenic activity). We purified and characterized the corresponding enzyme, namely NADPH-dependent 5β-dihydroprogesterone reductase, which is highly oxygen sensitive and whose corresponding genes are prevalent in *C. innocuum*. Moreover, *C. innocuum*–administered female C57BL/6 mice (aged 7 weeks) exhibited decreased serum progesterone levels (∼35%). *Clostridium*-specific antibiotics (metronidazole) restored low serum progesterone levels in these mice. Furthermore, prolonged *C. innocuum* administration (12 weeks) arrested ovarian follicular development in female mice. Cytological and histological analyses indicated that *C. innocuum* may cause luteal phase insufficiency and affect menstrual regularity. Our findings suggest *C. innocuum* as a causal factor of progesterone resistance in women taking progesterone.

## INTRODUCTION

Reproductive hormones, including progestogens (see **Figure S1** for the common structures of progestogens) and estrogens, modulate the metabolism, development, reproduction, and behavior of humans and other vertebrates^1^. Progesterone plays a crucial role in fertilization, implantation, and embryogenesis in women and female mice^2^. Progesterone has crucial functions in preparing the uterus for pregnancy and in maintaining pregnancy. Specifically, progesterone prepares the endometrium for implantation. In women, progesterone is mainly produced by the corpus luteum in the ovaries through two enzymatic steps; in the first step, the cholesterol side-chain cleavage enzyme (P450scc) transforms cholesterol into pregnenolone in the mitochondria, and in the second step, the human enzyme 3β-hydroxysteroid dehydrogenase converts pregnenolone into progesterone^3^.

Progesterone is primarily metabolized in the liver by the enzymes 5α-reductase and 3α-hydroxysteroid oxidoreductase, generating a series of metabolites [i.e., 3α-hydroxy-5α-pregnan-20-one (allopregnanolone)]. Allopregnanolone has protective and mood-stabilizing effects during pregnancy and the postpartum period^4,5^. Progesterone and its metabolites are highly lipophilic and easily pass through the blood–brain barrier. These compounds thus accumulate in the brain and are considered neurosteroids^6,7^. Previous studies suggested that progesterone-derived neurosteroids, including allopregnanolone, are produced exclusively using human enzymes^3^.

In women, progesterone is present only during the luteal phase of the menstrual cycle. During pregnancy, serum progesterone levels are in the nanomolar range^3^. Before childbirth, maternal progesterone levels may reach 200–2,000 nmol/L^8^. Insufficient progesterone levels can (*i*) lead to luteal phase deficiency, affecting fertility and increasing the risk of miscarriage, and (*ii*) affect embryo implantation and increase the risk of uterine endometrial pathologies^3^. For example, in women with chronic anovulation disorders such as polycystic ovary syndrome, insufficient progesterone levels are one of the most common causes of infertility and early miscarriage ^9–11^. Adequate progesterone supplementation in the luteal phase is essential to support successful embryo implantation and to prevent early miscarriage during fresh and frozen embryo transfer for women undergoing *in vitro* fertilization.

The bioavailability of supplemented progesterone varies greatly among individuals and is often quite low (approximately 10%)^12^. In the first-pass effect, progesterone metabolism by hepatic enzymes is a major mechanism of low bioavailability. However, more than half of a sample of progesterone may be metabolized extrahepatically^13^. Steroid hormones are recycled between the liver and gut through enterohepatic circulation. The reabsorption of steroids mainly occurs in the small intestine in humans^14^ and in the caecum in mice^15,16^. Recent studies have indicated the crucial role of gut microbes in the modification and reabsorption of androgens in the gut^17^. Similarly, circulating progesterone, derived from either endogenous production or exogenous administration, has been reported to be associated with the gut microbiota composition^18–20^. However, the microbial species and corresponding enzymes responsible for progesterone metabolism in the gut remain unknown.

*Clostridium innocuum* is a Gram-positive, spore-forming anaerobe that significantly affects the metabolism and functioning of the gastrointestinal tract in humans. A comparative genomic analysis of publicly available microbiota datasets related to healthy individuals indicated a high prevalence of *C. innocuum* in the global population (>80%)^21^. Despite being previously recognized as a commensal bacterium, *C. innocuum* has recently been associated with multiple health conditions, including diarrhea, bacteremia, endocarditis, osteomyelitis, and peritonitis^22–27^, This finding suggests that *C. innocuum* is an opportunistic pathogen; however, a related virulence mechanism is yet to be identified. In this present study, we identified *C. innocuum* as a major species responsible for gut progesterone metabolism. Moreover, we applied integrated omics approaches to identify the progestogenic metabolites, microbial enzymes, and corresponding genes. The oral administration of *C. innocuum* to female mice led to substantial decreases in serum progesterone levels and the arrest of both the estrous cycle and ovarian follicular development.

## RESULTS

### Anaerobic progesterone metabolism by the gut microbiota in female patients with infertility

Several studies have demonstrated that both endogenous progesterone and exogenous progesterone can alter the host gut microbiota. For example, progesterone treatment inhibited the germination of *Clostridium difficile* spores^28^ and led to the abundance of *Bifidobacterium*^19^ and *Lactobacillus*^29^ in the guts of female mice. Furthermore, progesterone has been reported to be metabolized by gut microbes. Kornman and Loesche^30^ demonstrated that *Bacteroides* species likely utilized progesterone for their growth. Therefore, we hypothesized that female patients with infertility may harbor progesterone-metabolizing microbes that utilize progesterone in the gut and inhibit the enterohepatic circulation of progesterone. To validate this hypothesis, we collected fresh fecal samples from 14 female patients with infertility (aged 20–40 years; see **Table S1** for detailed information) who received oral progesterone administration for endometrial preparation and thawed embryo transfer. Among these patients, Patient 1 exhibited characteristics symptoms of chronic anovulation, obesity, and considerably low bioavailability of oral progesterone supplementation (**Table S1**). Therefore, we investigated the progesterone metabolic activity of the gut microbiota from Patient 1. This patient’s fecal sample (approximately 0.5 g) was anaerobically incubated with progesterone (1 mM) in a chemically defined medium with mineral salts (DCB-1)^31^ or in Brain Heart Infusion (BHI) medium (a nutrient-rich medium). Progesterone-derived microbial products were extracted and identified through ultraperformance liquid chromatography (UPLC)–atmosphere pressure chemical ionization (APCI)–high-resolution mass spectrometry (HRMS). We used 10 progestogens (see **Figure S1** for the individual structures) as authentic standards; compared with progesterone, these standards are reduced at the 3-keto, 20-keto, and/or C-5 groups (the UPLC– HRMS manners of the individual progestogens are shown in **Table S2)**. In progesterone (1 mM)-amended fecal cultures, we observed apparent progesterone utilization by the gut microbiota, which was consistent with the marked decrease in progestogenic activity observed over time (**Figure 1Ai**).

**Figure 1.**
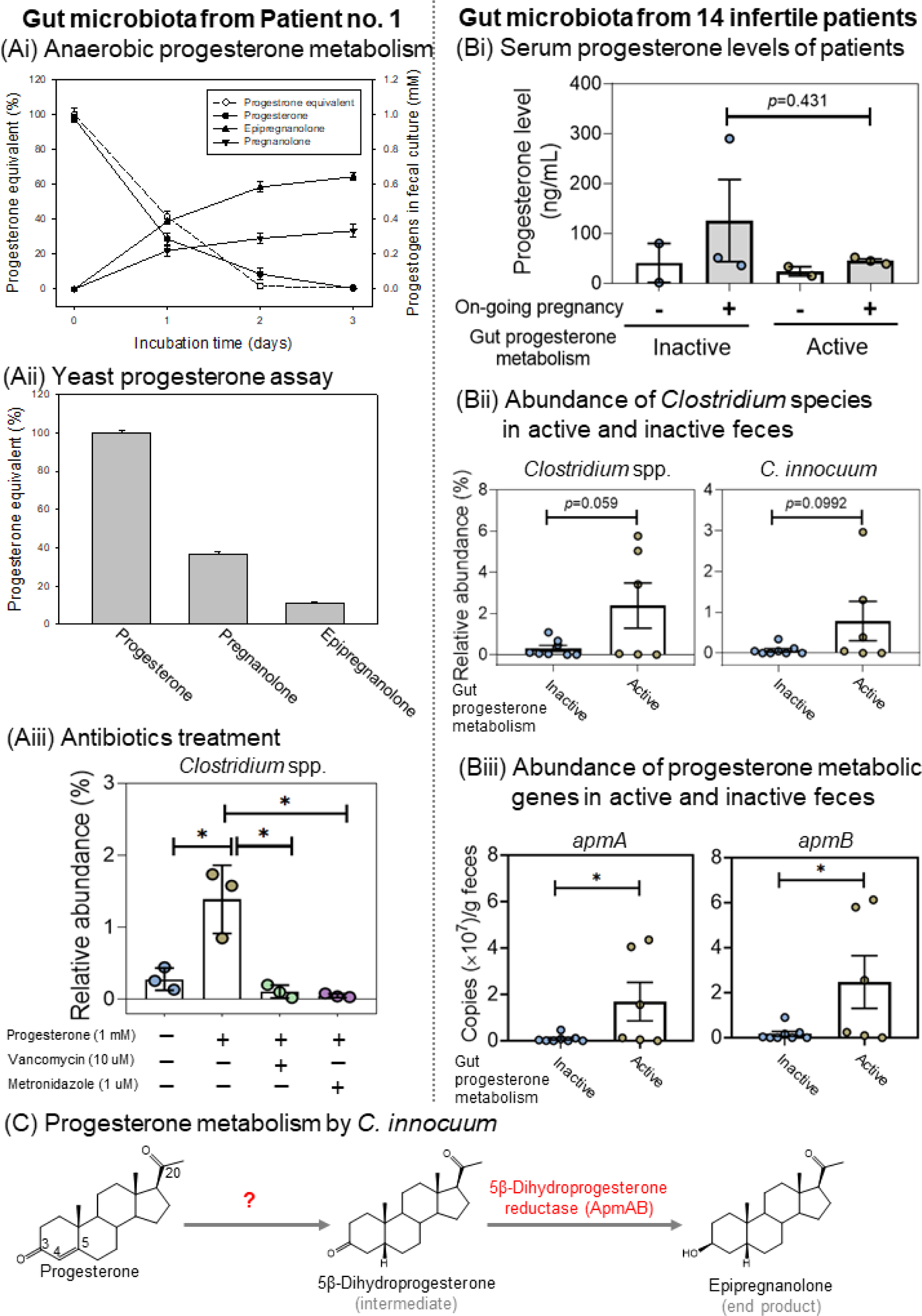
Identification of key players in gut progesterone metabolism and their corresponding genes. (**Ai**) Time-course progesterone utilization, microbial metabolite production, and progestogenic activities in fecal cultures. Progesterone-derived microbial products included 3β-hydroxy-5β-pregnan-20-one (epipregnanolone) and 3α-hydroxy-5β-pregnan-20-one (pregnanolone). (**Aii**) Relative progestogenic activities (%) of individual progestogens. Progestogenic activity was determined through the yeast progesterone assay. (**Aiii**) Inhibition of *Clostridium* growth and corresponding microbial activity by two *Clostridium*-specific antibiotics (see **Figure S3** for the effects of individual antibiotics on progesterone metabolism). The gut microbiota detailed in **Figures 1Ai–1Aiii** were sampled from a woman with infertility (Patient 1). Data are expressed as mean ± standard error for three independent experiments. (**B**) Differences in serum progesterone levels and abundance levels of progesterone-metabolizing gut microbes and their corresponding genes between patients exhibiting active and inactive gut progesterone metabolism. The gut microbiota were obtained from 14 women with infertility; gut microbiota from eight of these women exhibited negligible progesterone metabolic activity (Inactive group), whereas the gut microbiota from the remaining six women exhibited apparent progesterone metabolic activity (Active group). Detailed information regarding the physiological characteristics of 14 patients are provided in **Table S1**. (**Bi**) Serum progesterone levels of 10 patients with successful pregnancy in IVF were measured at the 3rd week after embryo transfer. Ongoing pregnancy was determined by the presence or absence of the gestational sac. (**Bii**) Relative abundance of the bacterial genus *Clostridium* and the *C. innocuum* species in gut microbiota. (**Biii**) Abundance of the progesterone metabolism genes *apmAB* in fecal samples. (**C**) Progesterone metabolism products and the characterized progesterone metabolism enzyme of *C. innocuum*.

Two progesterone-derived microbial products, 3β-hydroxy-5β-pregnan-20-one (epipregnanolone; the major product) and 3α-hydroxy-5β-pregnan-20-one (pregnanolone; the minor product), were observed in both the BHI (**Figure 1Ai**) and DCB-1 cultures (**Figure S2**). We then determined the progestogenic activities of individual progestogens (i.e., progesterone, epipregnanolone, and pregnanolone) through a yeast progesterone assay, and the aforementioned two microbial products exhibited considerably low progestogenic activity (**Figure 1Aii**). Among the two products, epipregnanolone exhibited the lowest progestogenic activity (a decrease of 89% compared with progesterone). These data indicated that the fecal sample from Patient 1 harbored highly active gut microbes that were able to metabolize progesterone (with a 3-keto group) into 3α- or 3β-hydroxyl structures (the microbial products and enzyme characterized in this study are provided in **Figure 1C**), leading to a decrease in progestogenic activity.

### Identification and characterization of key gut microbes responsible for progesterone metabolism

Subsequently, we identified major progesterone-metabolizing gut microbes. We used different antibiotics to selectively inhibit specific microbial populations in the gut microbiota, and we observed the effects of these antibiotics on anaerobic progesterone metabolism. First, we used two broad-spectrum antibiotics (tetracycline and thiamphenicol) to inhibit anaerobic progesterone metabolism. This microbial activity was inhibited by a relatively high concentration of tetracycline (minimum inhibition concentration = 50 μM) and thiamphenicol (minimum inhibition concentration = 25 μM) (**Figures S3A and S3B**). We then used two first-line antibiotics typically applied for *Clostridium* infection treatment (vancomycin and metronidazole) to inhibit anaerobic progesterone metabolism in progesterone-amended fecal cultures. Progesterone metabolism was highly inhibited by metronidazole (minimum inhibition concentration = 1.5 μM) and moderately inhibited by vancomycin (minimum inhibition concentration = 10 μM) (**Figures S3C and S3D**); these results indicated that *Clostridium* spp. likely are responsible for the anaerobic progesterone metabolism.

Sequencing on the PacBio platform also revealed an increase in the abundance of the genus *Clostridium* in the fecal cultures of Patient 1 after anaerobic incubation with progesterone (1 mM) for three days (**Figures 1Aiii and S4**), whereas *Clostridium* spp. were not enriched in the progesterone-amended cultures supplemented with the *Clostridium*-specific antibiotics vancomycin (10 μM) and metronidazole (1 μM) (**Figure 1Aiii**).

This study investigated whether *Clostridium* spp. are highly abundant in the gut microbiota of women with infertility who have high progesterone metabolic activity (see **Table S1** for demographic and hormonal characteristics). Negligible progestogenic metabolic activity was observed in the gut microbiota of eight women (Inactive group; n = 8), whereas progesterone metabolic activity was observed in the gut microbiota of six women (Active group; n = 6) (**Table S3**). Three weeks after embryo transfer, pregnant patients in the Active group exhibited lower serum progesterone levels (although not significant; *p* = 0.431) than those in the Inactive group (**Figure 1Bi**). After embryo transfer, the serum progesterone levels of pregnant patients in the Inactive group continuously increased, whereas the levels in the Active group decreased over time (**Figure S5**). We observed a significantly higher abundance of the genus *Clostridium* in the gut microbiota in the Active group (**Figure 1Bii**; left panel). Similarly, the abundance of *C. innocuum* was higher in the active gut microbiota in the Active group (**Figure 1Bii**; right panel). The difference in the abundance of *Clostridium difficile*, a common human pathogen, between the Active and Inactive groups was negligible (**Figure S6**). Thus, our data indicated the key role of *Clostridium* species, especially *C. innocuum*, in gut progesterone metabolism.

### Isolation and characterization of *Clostridium* species from the gut microbiota

We isolated *Clostridium* species from the gut microbiota of Patient 1. In total, we isolated and characterized 22 gut *Clostridium* strains belonging to seven different *Clostridium* species (**Table S4**). Of these 22 strains, only five (5/22; 23%) exhibited progesterone metabolic activity, and all of them were *C. innocuum* (**Table S4**). All these five *C. innocuum* strains produced epipregnanolone as the main product after anaerobic incubation with progesterone, whereas pregnanolone was produced only in trace amounts. Additionally, these *C. innocuum* strains were highly sensitive to metronidazole (with minimum inhibition concentration at 1.3 and 0.3 μg/mL in the BHI broth and BHI agar, respectively); by contrast, these strains were less sensitive to vancomycin, which was consistent with the results of the antibiotic inhibition trials (**Figures Aiii and S3**). Our data thus indicated the crucial role of *C. innocuum* in gut progesterone metabolism. Because the *C. innocuum* strain RGG8 exhibited the highest progesterone metabolic activity (**Table S4**), we used the strain RGG8 as the model microorganism for subsequent genomic and enzymatic characterization.

### Characterization of progesterone metabolism enzymes from *C. innocuum* strain RGG8

Given that the reduction of the 3-keto group in progesterone significantly decreased its activity (**Figure 1Aii**), we purified and characterized the microbial enzyme responsible for the reduction of 3-keto group. The resting cell assays revealed that the extracellular electron carrier methyl viologen and the strong reductant titanium citrate were required for the progesterone metabolism of *C. innocuum* (**Figure S7**), which is a typical characteristic of the enzymatic reactions of O_2_-labile [4Fe-4S] proteins^32,33^. Moreover, in the resting cell assays, the ATPase inhibitor did not reduce progesterone metabolic activity; this observation implies that progesterone metabolism does not require ATP-dependent uptake to transport progesterone into the cytoplasm and that the enzymes responsible for the reduction of the 3-keto group are likely located on the cell membrane.

Through chromatographic separation, we purified and characterized the *C. innocuum* enzyme responsible for the anaerobic reduction of the 3-keto group. The purified enzyme utilized 5β-dihydroprogesterone as the optimal substrate and did not use progestogens with a double bond at C-4 (*e.g.*, progesterone, 20α-dihydroprogesterone, and 20β-dihydroprogesterone) as the substrate (**Figure 2Ai**). With 5β-dihydroprogesterone serving as the substrate, the optimal pH of the purified protein was approximately pH 8.5 (**Figure S8A**), and the optimal working temperature was 25°C. Purified 5β-dihydroprogesterone reductase was still active (relative activity > 60%) at 45°C (**Figure S8B**). For 5β-dihydroprogesterone reductase, NADPH was the preferred electron donor (**Figure 2Aii**). Additionally, this enzyme was highly sensitive to O_2_. Under aerobic conditions, the activity of 5β-dihydroprogesterone reductase was partially maintained with the addition of a high concentration of the reductant 2-mercaptoethanol (**Figure 2Aiii**).

**Figure 2.**
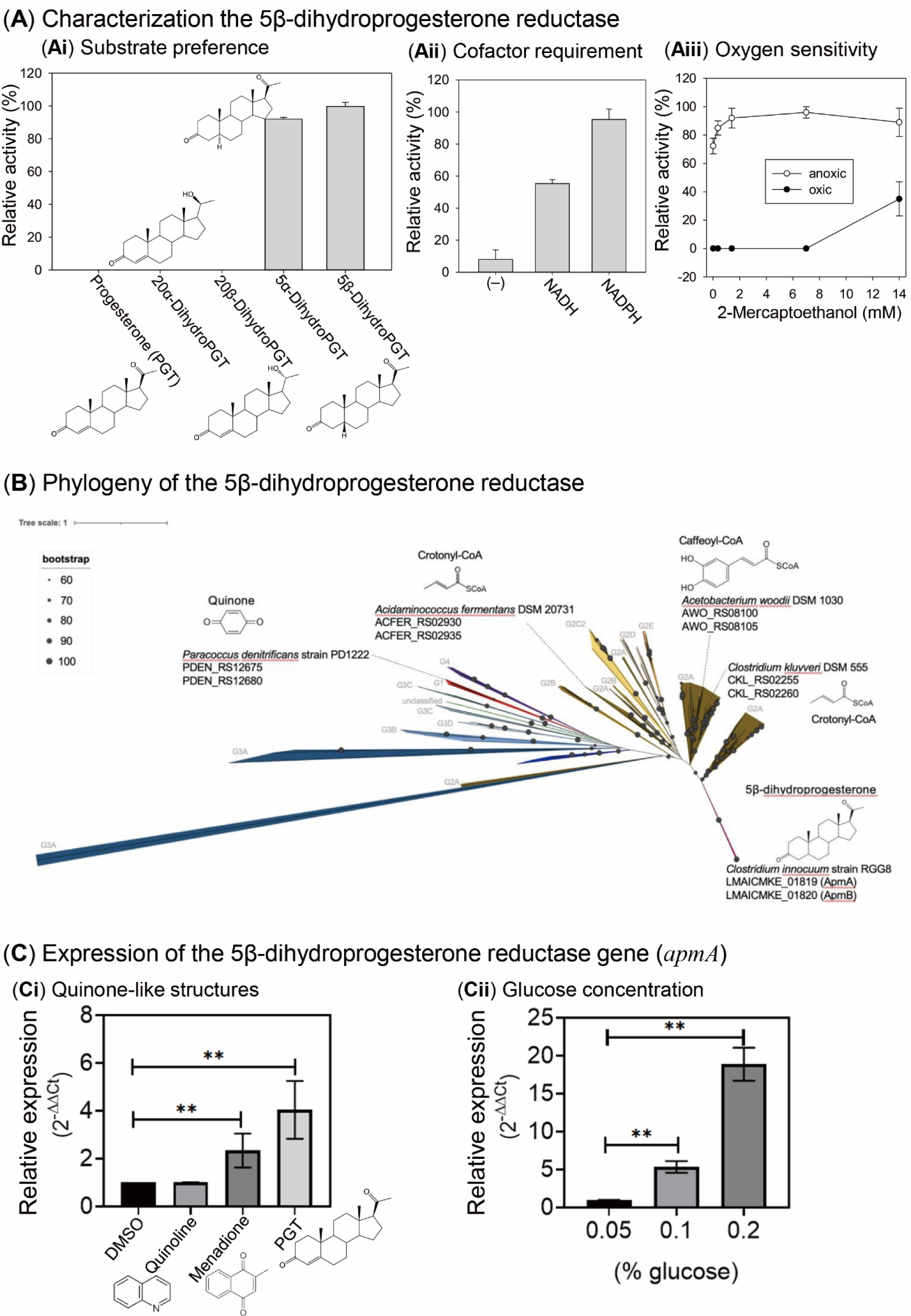
(**A**) Characterization of NADPH-dependent 5β-dihydroprogesterone reductase (ApmAB) purified from *C. innocuum* strain RGG8. (**B**) Phylogenetic analysis of 356 Etfαβ sequences, (including 206 ApmAB). Phylogenetic relationship was determined using Maximum-likelihood method and the bootstrap values were calculated based on 500 replicates. (**C**) Expression of *apmA* induced by progesterone (**Ci**) and glucose (**Cii**). All three cyclic compounds had a working concentration of 0.3 mM. The final DMSO concentration in the assay mixtures was 0.3% (v/v). Abbreviations: PGT, progesterone.

Proteomics analysis of the closed genome of the strain RGG8 revealed tryptic peptides originating from the active protein fraction match with the predicted tryptic products of strain RGG8 genes LMAICMKE_01819 (*apmA*) and LMAICMKE_01820 (*apmB*). Bioinformatic analysis indicated that ApmAB represented a new member of the electron-transferring flavoprotein (Etf) family, and the results suggested that both ApmA and ApmB contain the FAD-binding domain, whereas only the ApmA component possesses a [4Fe-4S] cluster as the prosthetic group. In addition to ApmAB in the strain RGG8, we collected 205 ApmAB sequences (identity > 98%) from publicly available *C. innocuum* genomes; using these data, we elucidated the phylogenetic relationship of ApmAB and other Etf members (150 sequences in total)^23^. We then constructed a phylogenetic tree with 356 concatenated Etfα and Etfβ protein sequences (**Figure 2B**). On the basis of this phylogenetic tree, we classified Etfαβ into five groups, which were similar to those reported by Costas et al.^23^. The ApmAB sequences formed a distinct subgroup within the G2 group. Noticeably, *apmAB* are prevalent in *C. innocuum* and they have not been identified in other *Clostridium* species (**Figure 2B**). Given that *apmAB* are highly conserved among *C. innocuum* species, we designed specific primers for *apmA* and *apmB*. Our studies regarding the expression of the *apmA* (the [4Fe-4S]-containing component) revealed that this gene can be induced by progesterone and menadione (another quinone-like structure), albeit to a lesser extent (**Figure 2Ci**). By contrast, the aromatic quinoline did not induce the expression of *apmA*. Contrary to our expectations, both *apmA* expression (**Figure 2Cii**) and the corresponding progesterone metabolic activity (**Figure S9**) were induced by glucose in a dose-dependent manner.

Subsequently, we determined whether *apmAB* were present in women with infertility who had gut microbiota exhibiting high progesterone metabolic activity. We extracted the total bacterial DNA from fresh fecal samples to determine the abundance of *apmA* and *apmB* in individual fecal samples. The quantitative polymerase chain reaction (PCR) results revealed a significantly higher abundance of both *apmA* (**Figure 1Biii**; left panel) and *apmB* (**Figure 1Biii**; right panel) in the gut microbiota with high progesterone metabolic activity, especially in the microbiota from Patients 1, 11, and 13 (**Table S3**). Together, these data support the crucial role of ApmAB from *C. innocuum* in anaerobic progesterone metabolism in the guts of patients with infertility, with epipregnanolone as the sole microbial product.

### Effect of *C. innocuum* on circulating progesterone levels, the estrous cycle, and follicle development in female mice

The present experiments suggested that *C. innocuum* may be a causal factor contributing to the low bioavailability of orally administered progesterone. We performed animal experiments using C57BL/6 mice as the animal model and investigated the effects of *C. innocuum* on the bioavailability of oral progesterone supplementation. Except for negative controls (without oral gavage; n = 5), all the tested mice were orally administered with progesterone (20 mg/kg/day). Female mice were administered with the strain RGG8 through oral gavage for 21 days (**Figure 3Ai**), and serum progesterone levels were subsequently determined through an enzyme-linked immunosorbent assay (ELISA). Compared with those of the negative control group (CTL; 9.7 ± 0.6 ng/mL), the serum progesterone levels of the progesterone-treated mice significantly increased (15.4 ± 1.3 ng/mL, n = 5). We used a common human pathogen, namely *C. tertium*^34^, as the reference *Clostridium* species (a negative control without progesterone metabolic activity). We did not observe any difference in serum progesterone levels among the mice treated with *C. tertium* (11.9 ± 1 ng/mL; n = 5), the mice treated with basal minimal medium (12.7 ± 1.5 ng/mL; n = 5), and the mice treated with progesterone alone. By contrast, serum progesterone levels (3.5 ± 0.2 ng/mL) significantly decreased in *C. innocuum*-treated mice (n = 5). However, metronidazole treatment recovered the *C. innocuum*– mediated decrease in serum progesterone levels (12.5 ± 0.7 ng/mL; n = 5) (**Figure 3Aii**). During the entire 21-day exogenous progesterone treatment period, we assessed the stage of the mouse estrous cycle through the daily cytological examination of vaginal smears. In addition, we recorded the body weight of each mouse on a weekly basis; however, we did not observe any changes in the stage of the mouse estrous cycle or body weight.

**Figure 3.**
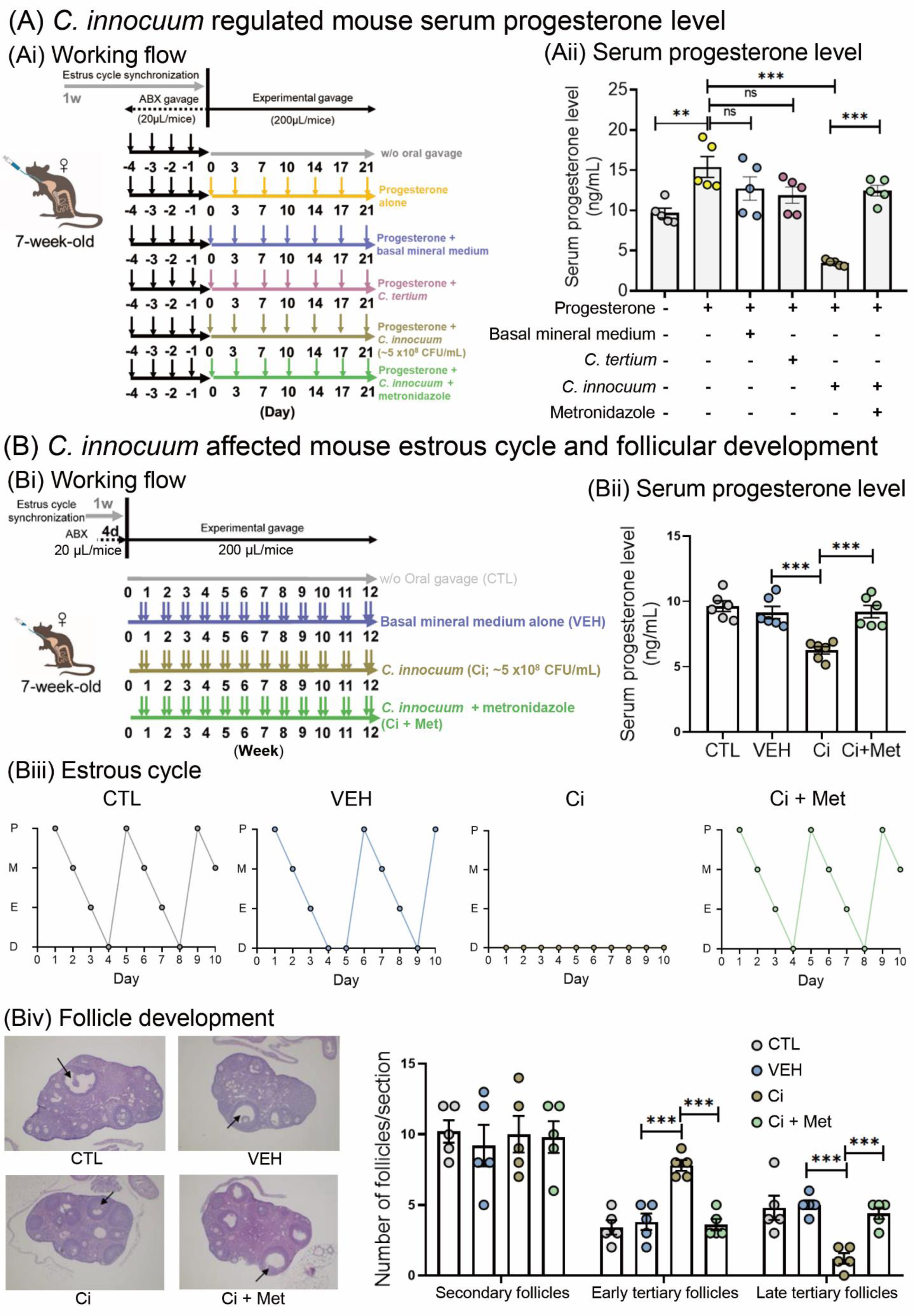
Administration of the progesterone-metabolizing *C. innocuum* strain RGG8 into the guts of female mice through oral gavage led to a decrease in host circulating progesterone levels and the arrest of ovarian follicular development. (**A**) Effects of *C. innocuum* on the bioavailability of oral progesterone supplementation. (**Ai**) Working flow of the administration of the strain RGG8 and progesterone in female mice through oral gavage for 21 days. (**Aii**) Administration of the strain RGG8 significantly reduced serum progesterone levels in female mice treated with progesterone. (**B**) Long-term effects of *C. innocuum* on host reproductive physiology. (**Bi**) Working flow of the administration of the strain RGG8 for 12 weeks. Prolonged administration of the strain RGG8 not only reduced serum progesterone levels (**Bii**) but also led to the arrest of the mouse estrous cycle (**Biii**) and follicular development (**Biv**). Data are expressed as mean ± standard error for six individual mice. Statistical results were calculated using the unpaired nonparametric *t*-test; \**p* < 0.05, \*\*\**p* < 0.001. Abbreviations: Ci, *C. innoccum* administration; Ci + Met, *C. innoccum* and metronidazole coadministration; CTL, female mice without oral gavage; VEH, vehicle mice.

To investigate the long-term effects of *C. innocuum* on host reproductive physiology, we administered *C. innocuum* strain RGG8 to female mice twice per week through oral gavage for 12 consecutive weeks (**Figure 3Bi**). Compared with the mice in the negative control group (CTL; without oral gavage; 9.6 ± 0.4 ng/mL; n = 6) and those in the vehicle group (VEH; with mineral medium; 9.1 ± 0.5 ng/mL; n = 6), serum progesterone levels were significantly decreased in the *C. innocuum*-treated mice (6.3 ± 0.3 ng/mL; n = 6). Coadministration with metronidazole restored the decline in serum progesterone levels (9.2 ± 0.5 ng/mL; n = 6) (**Figure 3Bii**). In addition, prolonged *C. innocuum* administration led to a decrease in serum estradiol levels but increases in the serum levels of follicle-stimulating hormone (FSH) and luteinizing hormone (LH) (**Figure S11**). The original levels of these hormones were restored by coadministration with metronidazole. Moreover, cytological analysis of vaginal smears demonstrated that the 12-week *C. innocuum* administration led to an arrest of the mouse estrous cycle, which was then restored by metronidazole treatment (**Figure 3Biii**). Furthermore, the histological analysis of the ovarian tissue revealed that compared with the CTL and VEH groups, *C. innocuum* administration significantly increased the number of early tertiary follicles but decreased the number of late tertiary follicles. Overall, our data suggest that prolonged *C. innocuum* administration leads to ovarian follicle arrest in the early antral stage but that such arrest can be eliminated by coadministration with metronidazole (**Figure 3Biv**).

## DISCUSSION

To the best of our knowledge, the present study is the first study to provide robust evidence demonstrating that gut progesterone serves as an electron acceptor for the strictly anaerobic *C. innocuum* and that the respiration process occurs extracellularly and does not require progesterone uptake. Gut microbiota provide a highly anaerobic and reductive environment^35,36^ with abundant electron donors but few electron acceptors. Similar to quinones, the A-ring of progesterone has conjugated double bonds at C-3 and C-5, which can accept two pairs of electrons. In this study, we purified and characterized NADPH-dependent 5β-dihydroprogesterone reductase, which belongs to the Etf family. Moreover, bioinformatic analysis of 5β-dihydroprogesterone reductase (ApmAB) from *C. innocuum* revealed that this enzyme is similar to the CarDE component of caffeoyl-CoA reductase from *Acetobacterium woodii* (amino acid sequence identity = approximately 50%). In both subunits of heterodimeric 5β-dihydroprogesterone reductase, we identified a highly conserved Rossman domain for binding NADPH. Electrons are likely transferred sequentially from NADPH, FAD, and finally to progesterone, with epipregnanolone as the main product.

In a previous study, 3β-hydroxysteroid dehydrogenase was considered an alternative enzyme capable of reducing the 3-keto group in various steroids, including progesterone ^37^. The protein 3β-hydroxysteroid dehydrogenase [belonging to the short-chain dehydrogenase/reductase (SDR) family] is widely distributed in yeast^38^, plants^39^, animals^40^, and bacteria^41,42^. These proteins possess a highly conserved NAD^+^-binding domain and are not oxygen sensitive^39^. In one study, NAD^+^-dependent 3β-hydroxysteroid dehydrogenase was purified from the cytoplasm of *C. innocuum*^37^. This monomeric enzyme was able to mediate the reverse reaction (reduction of the 3-ketosteroids into 3β-hydroxysteroids), in which NADH was used as electron donor. However, the enzymatic activity is very low (specific activity = 16 nmol progesterone consumed/hour/mg protein) and was not physiologically relevant. Moreover, the oxygen tolerance of the SDR-type 3β-hydroxysteroid dehydrogenase was inconsistent with the results of the present resting cell assay, which revealed the oxygen sensitivity of progesterone-metabolizing enzymes in *C. innocuum*. Accordingly, in *C. innocuum*, NAD^+^-dependent 3β-hydroxysteroid dehydrogenase may catalyze the oxidation of 3β-hydroxysteroids in the cytoplasm, and 5β-dihydroprogesterone reductase (ApmAB) may catalyze the 3-keto reduction of 3-ketosteroids on the cell membrane.

Steroids with pregnane skeleton are effective modulators of neuronal γ-aminobutyric acid (GABA) receptors^43^. The spatial arrangement of the hydroxyl group at the C-3 position of the steroid skeleton markedly affects the biological properties of steroids. Pregnanolone and epipregnanolone are stereoisomers of allopregnanolone, and they differ from allopregnanolone in terms of the orientations of the H-5 and 3-hydroxy groups. 3α-Hydroxypregnanes (e.g., allopregnanolone) are strong positive modulators, and in nanomolar concentrations, 3α-hydroxypregnanes can potentiate the GABA-induced chloride current; 3β-hydroxypregnanes (e.g., epipregnanolone) exhibit antagonistic properties and weaken the stimulating effect of allopregnanolone on the GABA-induced chloride current^44^. Allopregnanolone is considered as a neuroprotective agent^4,5,45^. Other neurosteroids have not yet attracted intensive research attention. Some studies have demonstrated that epipregnanolone may have therapeutic effects that extend beyond the modulation of GABA receptors. For example, O’Dell et al. (2005)^46^ described a significant reduction of alcohol self-administration following pre-treatment with epipregnanolone, and they proposed that this effect may be related to epipregnanolone’s actions on the GABA receptor. However, contrasting results have been obtained regarding the effects of 3-hydroxypregnanes on GABA receptors, and some investigators have described epipregnanolone as a positive modulator of GABA Receptor^47^.

In the present study, we observed that patients with high progesterone metabolic activity in their fecal samples exhibited low circulating progesterone levels, and we also observed the unusual capability of *C. innocuum* to decrease circulating progesterone levels, likely through the inactivation of endogenous and exogenous progesterone in the mouse gut. Thus, we hypothesize that *C. innocuum* is a causal factor of progesterone resistance in women with infertility who take progesterone. Accordingly, individualized progesterone dosage could be determined based on the progesterone metabolism capability of the gut microbiota of a patient; such determination could not only reduce costs but also minimize the side effects associated with high-dose medications, thereby enhancing treatment efficiency. This study also observed the arrest of the follicular development and estrous cycle in mice with prolonged *C. innocuum* administration. In humans, progesterone is mainly produced by the corpus luteum in the ovaries^48^. Follicular arrest may lead to the dysfunction of the corpus luteum, which further hinders endogenous progesterone production. Therefore, microbial activities and host follicular arrest (through an unidentified route) may lead to the lower progesterone levels in both female mice and women with infertility who have high *C. innocuum* abundance. In addition to progesterone, prolonged *C. innocuum* administration led to marked changes in serum levels of estradiol, progesterone, FSH, and LH in this study. These findings imply a potential role of *C. innocuum* in human ovulatory disorders. Nonetheless, the effects of *C. innocuum* on pregnancy, miscarriage risk, and related disorders resulting from progesterone metabolic abnormalities remain to be investigated through clinical trials with a larger sample size.

## Conclusions

In this study, we identified *C. innocuum* as a major microbe that utilizes progesterone in the gut. This microbial process occurring in the gut can significantly decrease the host circulating progesterone levels through enterohepatic circulation. The biotransformation of progesterone into epipregnanolone in the host gut hinders not only the progestogenic activity of progesterone but also intestinal progestogen reabsorption, resulting in decreased circulating progesterone levels in the host. Our findings provide promising therapeutic targets for the clinical amelioration of low bioavailability of progesterone. For example, treatment with antibiotics (e.g., *Clostridium*-specific metronidazole) can be used to eliminate the gut *C. innocuum* population before oral progesterone supplementation.

Moreover, the genes corresponding to 5β-dihydroprogesterone reductase from *C. innocuum* are prevalently in *C. innocuum* but not identified in any other *Clostridium* species. Thus, these highly specific functional genes, especially *apmA*, can be used as clinical biomarker for identify patients with infertility who have low serum progesterone levels and low bioavailability of oral progesterone supplementation. To the best of our knowledge, this study is the first study to provide strong evidence demonstrating that gut microbes can produce neurosteroids from progesterone.

Progesterone-derived compounds with 3-hydroxyl groups are potential neurosteroids^3^. Currently, most progesterone-derived neurosteroids are either extremely expensive or not available commercially, which considerably impedes detailed investigations of the physiological and psychological effects as well as the clinical applications of these neurosteroids. Compared with the production of neurosteroids through organic chemical approaches, the use of gut microbes (e.g., *C. innocuum*) and their enzymes may result in the production of neurosteroids with relatively few stereoisomers and regioisomers, with epipregnanolone, for example, being the sole product; progesterone-transforming enzymes with high regio- and stereo-specificity thus have many biotechnological applications. Neurosteroids production through microbial or enzymatic processes could considerably decrease the cost of chromatographic purification.

## MATERIALS AND METHODS

Human fecal samples (approximately 0.5 g each) were anaerobically incubated with progesterone (1 mM) in DCB-1 medium or BHI broth. Bacterial DNA was extracted and amplified through PCR, and the bacterial 16S rRNA amplicons were sequenced on a PacBio platform. The abundance of progesterone metabolism genes in fecal samples was determined through quantitative PCR (qPCR). Progesterone-derived microbial products were identified through UPLC–APCI– HRMS, and their progestogenic activity [presented as the progesterone equivalent (%)] was determined through a yeast progesterone assay. Progesterone-metabolizing bacteria, including the *C. innocuum* strain RGG8, were isolated from human feces through a culturomics approach. The strain RGG8 was routinely grown in BHI broth containing progesterone (1 mM) under strictly anaerobic conditions. The genome of the strain RGG8 was sequenced on the PacBio platforms. The expression of the functional genes of strain RGG8 was determined through reverse transcription (RT)–qPCR. Bacterial cells were destroyed using a French Press in an O_2_-free system. Protein purification was performed on a fast protein liquid chromatography (FPLC) system in an anaerobic chamber.

Bacterial proteins were separated sequentially through ammonium sulfate precipitation and by using columns separately containing DEAE Sepharose, Phenyl Sepharose, and Sephacryl S-300. The active protein fraction was treated with trypsin, and the tryptic peptides were analyzed through UPLC–MS. The strain RGG8 was administered to the mouse gut through oral gavage. Healthy C57BL/6 female mice (aged 7 weeks) were used as model hosts. The C57BL/6 mice were obtained from the Animal Center of the Medical College (National Taiwan University, Taipei, Taiwan) and were maintained under standard animal housing conditions according to the Guide for the Care and Use of Laboratory Animals. Serum hormone levels of mice were measured using ELISA. Progestogenic metabolites in fecal and serum samples were extracted using ethyl acetate and were then analyzed through UPLC– HRMS. The mouse estrous cycle and follicle development were investigated through cytological and histological analyses, respectively. The materials and methods of this study are described in further detail in the Supporting Information.

## Acknowledgements

We thank the Small Molecule Metabolomics core facility sponsored by the Institute of Plant and Microbial Biology, Academia Sinica (Taiwan), for its UPLC–HRMS analysis (AS-CFII-111-218).

## Author contributions

M.-J.C., C.-H.C., T.-H.H., P.-H.W., and Y.-R.C. designed the research; C.-H.C., T.-H.H., T.-Y.W., C.-Y.L., Y.-L.C., R.G.G., G.-J.B.-M., Y.-L.L, P.-T.L., and Y.-L.T. conducted the research; P.-H.W., T.-H.L., and C.-J.S. contributed new reagents and analytic tools; M.-J.C., C.-H.C., T.-H.H., P.-H.W., and Y.-R.C. analyzed the data; and M.-J.C., C.-H.C., T.-H.H., P.-H.W., and Y.-R.C. composed the manuscript.

## Data availability

Genomic data related to the strain RGG8 are available in **Dataset S1**. Genes of the Etf protein family selected for phylogenetic analysis are detailed in **Dataset S2**.

## Ethics approval

Human fecal sample collection was approved by the Clinical Trial /Research Approval NTUH-REC No.:202103046RINB). This animal study conducted in accordance with the requirements of the Taiwan Animal Protection Law on Scientific Application of Animals and the Institutional Animal Care and Use Committee (IACUC) of National Taiwan University College of Medicine (IACUC No. 20220423).

