## Supplemental Information for "*Clostridium innocuum*, an opportunistic gut pathogen, inactivates host gut progesterone and arrests ovarian follicular development"

**This PDF file includes:**

Supplemental Materials and Methods
SI References
Figures S1 to S11
Table S1 to S5
Legends for Datasets S1 and S2

**Other supplementary materials for this manuscript include the following:**

Dataset S1
Dataset S2

### Supplemental Materials and Methods

#### Chemicals

Progesterone was obtained from Sigma-Aldrich (St. Louis, MO, USA). The other progestogens, including 5α-dihydroprogesterone, 5β-dihydroprogesterone, 3α-hydroxy-5α-pregnan-20-one (allopregnanolone), 3β-hydroxy-5α-pregnan-20-one (isopregnanolone), 3β-hydroxy-pregnan-20-one, 3α-hydroxy-5β-pregnan-20-one (pregnanolone), 3β-hydroxy-5β-pregnan-20-one (epipregnanolone), 3α-hydroxy-4-pregnen-20-one, 3β-hydroxy-4-pregnen-20-one, 20α- dihydroprogesterone, and 20β- dihydroprogesterone were purchased from Steraloids (Newport, Rhode Island, USA). Other chemicals used were of analytical grade and were purchased from Mallinckrodt Baker (Phillipsburg, USA), Merck Millipore (Burlington, USA), and Sigma-Aldrich unless specified otherwise.

#### Identification of microbial progestogenic metabolites and determination of progestogenic activity in the fecal cultures

Fresh fecal samples were collected from 14 infertile females (aged 20~40 years; see **Table S1** for the detailed information) who received oral progesterone administration for endometrial preparation and thaw embryo transfer. The fecal samples (approximately 0.5 g for each) was anaerobically incubated with progesterone (1 mM) in a chemically defined mineral salts medium (DCB-1; 100 mL)[31] or Brain Heart Infusion medium (BHI; 100 mL) at 37°C in the dark. 17α-Ethinylestradiol (final concentration = 50 μM), which cannot be utilized by gut microbiota, was added to fecal cultures to serve as an internal control. Cultures samples (3 mL) were withdrawn from the progesterone-amended fecal cultures daily. Progesterone-derived microbial products were then extracted from cultural samples (1 mL) with ethyl acetate twice After the solvent was completely evaporated, the residue was re-dissolved in 10 μL of methanol and progestogenic metabolites were identified using thin-layer chromatography (TLC) and ultraperformance liquid chromatography–atmosphere pressure chemical ionization–high-resolution mass spectrometry (UPLC–APCI–HRMS). We used 10 progestogens (see **Figure S1** for individual structures) as authentic standards, which are reduced at the 3-keto, 20-keto, and/or C-5 groups (the UPLC–HRMS manners of individual progestogens are shown in **Table S2)**. To determine the progestogenic activity in the fecal cultural samples, the samples (1 mL) were extracted with ethyl acetate three times; after the solvent was completely evaporated, the residue was re-dissolved in dd$H_2O$ to determine its progestogenic activity (see below). To determine temporal changes in bacterial community structures of the fecal cultures, bacterial cells were collected through centrifugation (10,000 × *g* for 10 min; 4 °C) and the Bacterial DNA was extracted from the pellet using QIAamp® PowerFecal® Pro DNA kit (Qiagen, Hilden, Germany) according to the manufacturer's instructions. The DNA concentration was determined using NanoDrop Spectrophotometer ND-1000 or Qubit™ dsDNA Assay kit (Invitrogen Thermo Fisher Scientific, Waltham, MA, USA). The bacterial 16S rRNA was amplified through PCR and the resulting amplicons were sequenced on a PacBio platform (see below). To investigate the effects of antibiotics on microbial progesterone metabolism, the fecal sample (approximately 0.5 g) collected from Patient no. 1 was anaerobically incubated with progesterone (1 mM) and individual antibiotics in the BHI broth (1 mL) at 37°C in the dark. Samples (3 mL) were withdrawn from the progesterone-amended fecal cultures daily.

#### Culturomics approach for isolation of progesterone-metabolizing microbiota

Progesterone-metabolizing bacteria were isolated from human fecal samples using a culturomics approach. In brief, fecal samples (5 g) were suspended in anaerobic DCB-1 broth (100 mL) and shaken vigorously. The resulting suspensions were serially diluted (up to $10^{-6}$), and 100 uL aliquot of the $10^{-3}$ to $10^{-6}$ dilutions were plated on agars specific for different bacterial taxa. The agars were incubated in the anaerobic chamber (Coy Laboratory Products, MI, USA). Amon them, Gifu Anaerobic Medium (GAM) (BD-Difco, MD, USA) was used for the general isolation of gut anaerobes, including those belonging to the Family Clostridiales. *Clostridium butyricum* Isolation Medium (BIM) (Popoff, 1984) and Iron Sulfite Agar (Hi-Media, India) were used for the isolation of *C. butyricum* and the related species. *Bacteroides Fragilis* Selective Medium (BFS) (Ho et al., 2017) was used to isolate the members of Bacteroidetes. *Raffinose-Bifidobacterium* (RB) Medium (Hartemink et al., 1996) and the BFM Selective Medium (Nebra and Blanch, 1999) were used to isolate the Bifidobacteria and related toxon, and De Man, Rogosa, and Sharpe (MRS) Medium (BD-Difco, MD, USA) was used to isolate the Lactic Acid Bacteria (LAB) and the related taxon. Progesterone (1 mM) was added into all growth media. The anaerobic growth media were prepared using the Hungate station; the bacterial growth on agar were performed in the anaerobic chamber. All glassware, media, petri dishes, and reagents were kept anaerobically. After anaerobic incubation for 2 to 5 days at 37°C, colonies were observed, and individual colonies were picked and inoculated to 800 uL GAM broth (using the 48-well plates). The 48-well plates were incubated at 37°C in an anaerobic chamber with shaking (100rpm) for 3 days. These active cultures (with $OD_{600nm} > 0.1$) were used as the pre-cultures for the high-throughput screening as described below.

**High-throughput screening for anaerobic progesterone metabolic activity**

Bacterial colonies were screened for anaerobic progesterone metabolic activity by transferring 100 ul of the pre-cultures to a fresh GAM broth in 24-well plates containing 1 mM of progesterone. The plates were incubated at 37°C with shaking (100rpm) for 24~48 h in an anaerobic chamber. After that, microbial activity was monitored using the High-throughput TLC analysis.  The active bacterial isolates were kept in GAM broth containing 20% glycerol or 5% DMSO and the resulting axenic cultures were stored at -80°C.

**Anaerobic growth of strain RGG8 on progesterone**

Strain RGG8 was routinely grown in the BHI broth (100 mL) containing progesterone (1 mM). 17α-Ethinylestradiol (50 µM; indigestible by strain RGG8) was added as an internal control. The anaerobic cultures were incubated in the dark at 37°C with stirring (approximately 180 rpm). The cultural samples (3 mL) were withdrawn every 12 h. The progesterone-derived metabolites extracted from the cultural samples were identified and quantified using UPLC–HRMS and the progestogenic activity in the cultural samples was determined using the yeast progestogenic activity assay as described below. To determine the bacterial growth (represented as the increase in protein content), strain RGG8 cultural samples were centrifuged at 10,000 × *g* for 10 min. After centrifugation, the cell pellet was resuspended in 1 mL of reaction reagent (Pierce BCA protein assay kit; Thermo Scientific). The protein content was determined using a BCA protein assay according to manufacturer’s instructions with bovine serum albumin as the standard.

**BioVAL A-YPS yeast progestogenic activity assay**

The yeast progestogenic activity assay was accomplished using the BioVAL A-YPS kit according to manufacturer’s protocol. In brief, the A-YPS kit uses the non-conventional and recombinant yeast *Arxula adeninivorans* as the progesterone responsive biosensor and the reporter protein, phytase, is directly secreted into the medium. There is a positive correlation between the phytase production and the progestogenic activity; the amount of phytase is thus able to be measured through spectrophotometrically monitored the enzymatic reaction using a chromogenic substrate. The individual progestogenic standards or the dried ethyl acetate extracts were dissolved in and diluted using dd$H_2O$. The resulting aqueous solutions (400 μL) were added to yeast cultures (100 μL; initial $OD_{600nm}$ = 0.5) in a 96-well microtiter plate and the bioassay mixtures were incubated at 30 ˚C for 24 h. Subsequently, the Substrate Buffer (50 μL) in the kit was added to the 96-well microtiter plate and the mixture was further incubated at 37 ˚C for 30 min. The reaction was then stopped by adding 100 μL of Developer solution into the bioassay mixture, the absorption of the bioassay mixtures was measured at a wavelength of 405 nm on a microtiter plate spectrophotometer.

**Analytical chemical methods**

**(a) Thin-layer Chromatography (TLC)**

The progestogenic standards and ethyl acetate-extractable compounds were separated on a silica gel-coated aluminum HPTLC plate (Silica gel 60 F254, thickness, 0.2 mm; 20 × 20 cm; Merck). The mobile phase was dichloromethane:ethyl acetate:ethanol (14:4:0.05, v/v/v). The compounds were visualized using UV light at 254 nm or by spraying the TLC plates with 30% (v/v) $H_2SO_4$, followed by heating on a hot plate.

**(b) Ultra-Performance Liquid Chromatography–High-Resolution Mass Spectrometry (UPLC–HRMS)**

Progestogenic metabolites were detected using UPLC–HRMS on a UPLC system coupled to an Atmosphere Pressure Chemical Ionization–Mass Spectrometry (APCI–MS) system. Progestogenic metabolites were separated using a reversed-phase $C_{18}$ column (Acquity UPLC® BEH $C_{18}$; 1.7 μm; 100 × 2.1 mm; Waters) at a flow rate of 0.45 mL/min at 50°C (oven temperature). The mobile phase was composed of a mixture of two solutions: solution A [0.1% formic acid (v/v) in 2% acetonitrile (v/v)] and solution B [0.1% formic acid (v/v) in acetonitrile (v/v)]. Separation was conducted by gradually increasing the concentration of solvent B from 40% to 55% in 9 min, then ramping up to 100% B within 1 min. Mass spectrometric data in positive ionization mode (parent scan range: 100–500 m/z) were collected. The temperatures of the capillary and APCI vaporizer were 120°C and 395°C, respectively; the sheath, auxiliary, and sweep gas flow rates were 40, 5, and 2 arbitrary units, respectively. The source voltage was 6 kV and the current was 15 μA. The elemental composition of individual adduct ions was predicted using Xcalibur™ Software (Thermo Fisher Scientific).

**Preparation of cell extracts of strain RGG8**

*C. innocuum* strain RGG8 was anaerobically cultivated in BHI broth (totally 10 L) containing 50 μM of progesterone as an inducer; the bacterial cells were harvested at $OD_{600nm}$ = approximately 0.8 through centrifugation. Cell pellet (10 g) was re-suspended in 30 mL suspension buffer (pH8.5) containing Tris-HCl (50 mM), 2-mercaptoethanol (10 mM), cOmplete™ Mini Protease Inhibitor, 5% glycerol, and 1 mM DNaseI. Strain RGG8 cells were broken by passing the cell suspensions through a French pressure cell (Thermo Fisher Scientific) twice at 137 megapascals in an $O_2$-free system. The cell-free lysates were fractionated using two steps of centrifugation steps. First, the cell-free lysates were centrifuged at 20,000 × *g* for 30 min to get rid of most cell debris and unbroken cells. Second, the supernatants containing the crude cell extracts were centrifuged at 100,000 × *g* for 1.5 h to fractionate the soluble proteins from the membrane-bound proteins. Before ultra-centrifugation, the cell-free lysate was treated with 0.2% (w/v) of the detergent Tween 20 to solubilize peripheral membrane proteins. All steps used for preparation of cell-extracts were performed at 4°C.

**Purification of the ApmAB from the crude cell extract of strain RGG8**

The crude cell extract was first precipitated at 30% ammonium sulfate saturation. After centrifugation, the pellet was re-dissolve in buffer A (50 mM Tris–HCl and 10 mM 2-mercaptoethanol; pH 8.5). The salts were removed using a prepacked PD-10 desalting column (8.3 mL; Amersham Biosciences Europa) according to the manufacturer's instructions. Further protein purification was performed on a Fast-Performance Liquid Chromatography (FPLC) established in an anaerobic chamber, and the bacterial proteins were separated sequentially through diethylaminoethyl (DEAE) Sepharose (ion exchange), Phenyl Sepharose (hydrophobic interaction), and Sephacryl S-300 (gel filtration). In brief, the bacterial proteins were applied to a DEAE Sepharose column (5 mL) previously equilibrated with buffer A (50 mM Tris–HCl, pH 8.5). The elution was performed using buffer B (50 mM Tris–HCl, 1 M KCl, and 10 mM 2-mercaptoethanol; pH 8.5) with a gradient from 0% to 50% within 30 min at a flow rate of 1 mL/min. The ApmAB activity was determined as described below. The active protein fractions were collected and then applied to a Phenyl Sepharose column (1 mL; GE Healthcare) equilibrated with buffer B. The elution was performed using buffer A with a gradient from 0% to 100% within 1 h at a flow rate of 0.5 mL/min. Active fractions were pooled and then concentrated using Vivaspin 500 centrifugal concentrator. The resulting proteins were applied to TSKgel® G3000SW HPLC Column equilibrated with buffer A (50 mM Tris–HCl, 100 mM KCl, and 10 mM 2-mercaptoethanol; pH 8.5) at a flow rate of 0.5 mL/min. Active fractions were pooled and then concentrated using a Vivaspin 500 centrifugal concentrator.

**Resting cell and *in vitro* enzymatic assays**

The resting cell assay and *in vitro* enzymatic assay were routinely performed in the dark at 30 °C under oxic or anoxic condition for 1.5 h. For the resting cell assay, *C. innocuum* strain RGG8 and *Eubacterium limosum* (as a reference strain) cells were harvested during the middle log phase (with an $OD_{600nm}$ = approximately 0.8) through centrifugation and the pellets were washed using Tris-HCl buffer (50 mM; pH 8.5) twice. The assay mixtures (0.5 mL) contained Tris-HCl buffer (50 mM; pH 8.5), cell pellet (0.1 g), progesterone (0.2 mM) and 2-mercaptoethanol (10 mM). In some assays, Ti(III) citrate (3 mM), methyl viologen (0.5 mM), and/or the ATPase inhibitor (sodium orthovanadate; 10 mM) were added to the assay mixtures. For the *in vitro* enzymatic assay, the assay mixtures (0.5 mL) contained Tris-HCl buffer (50 mM, pH 8.5), crude cell extract (for protein fractions) of strain RGG8, 5β-dihydroprogesterone (the substrate for ApmAB; 0.1 mM), NADPH (20 mM), and 2-mercaptoethanol (10 mM). After 1.5 hours of anaerobic incubation, the assay mixtures were extracted twice with the same volume of ethyl acetate. The ethyl acetate extracts were pooled, vacuum-dried, and stored at -20°C before further analytical analysis. TLC and UPLC–HRMS were used to monitor the epipregnanolone production.

**General molecular biological methods**

Bacterial genomic DNA, including that of strain RGG8, was extracted using the Presto™ Mini gDNA Bacteria Kit (Geneaid, New Taipei City, Taiwan). PCR mixtures (50 µL) contained nuclease-free $H_2O$, 2 × PCR master mix (Invitrogen™ Platinum™ Hot Start PCR 2X Master Mix, Thermo Fisher Scientific, Waltham, MA, USA), forward and reverse primers (200 nM each), and template DNA (10-30 ng). The PCR products were verified using standard TAE-agarose gel (1.5%) electrophoresis with the SYBR® Green I nucleic acid gel stain (Invitrogen Thermo Fisher Scientific, Waltham, MA, USA), and the PCR products were purified using the GenepHlow Gel/PCR Kit (Geneaid, New Taipei City, Taiwan). The TA cloning was performed with T&A™ Cloning Vector Kit (YEASTERN BIOTECH, New Taipei City, Taiwan).

**PacBio sequencing of the strain RGG8 genome**

For library preparation and PacBio sequencing, approximately 1 μg of strain RGG8 genomic DNA was sheared by Covaris g-TUBE (Covaris, Woburn, MA, USA) and purified via AMPure PB beads (PacBio, Menlo Park, CA, USA). The sheared and purified DNA fragments were used as templates to prepare the SMRTbell library through SMRTbell template prep kit 1.0 (PacBio, Menlo Park, CA, USA), according to the manufacturer's instructions. After adaptor-ligation to the inserts, the inserts with suitable size for sequencing were selected with the BluePippin system. The SMRT sequencing was performed on SMRT 1M Cell v3 (PacBio, Menlo Park, CA, USA) with chemistry version 3.0 on PacBio Sequel sequencer (Genomics BioSci & Tech Co). A primary filtering analysis was achieved on the Sequel System, and the secondary analysis was completed using the SMRT analysis pipeline version 8.0. For the genome assembly, the filtered subreads after SMRT Link v8.0 were assembled by a long-read assembly algorithm Flye v2.7. The SSPACE-LongRead v1.1 was applied for draft genome scaffolding and PBJelly v15.8.24 was used for gap closure. Subsequently, genome polishing was conducted with Arrow v2.3.3 software (PacBio, Menlo Park, CA, USA). A further assembly of the circularizing genome was conducted using Circlator v1.5.5. Finally, the quality of the assembled genome was evaluated by QUAST v4.5. After the *de novo* genome assembly, the annotations of genomic bacterial features were achieved with Prokka v1.13.

**Phylogenetic analysis of the ApmAB**

To elucidate the phylogenetic relationships of the 5β-dihydroprogesterone reductase (ApmAB) with other Etfαβ sequences, we constructed a maximum-likelihood (ML) phylogenetic tree using 356 concatenated Etfαβ-like sequences. These sequences originated from two primary sources: 205 ApmAB-like sequences derived from the publicly available genomes of *C. innocuum* and 150 Etfαβ sequences representative of five distinct groups, as identified in a foundational study on the systematic classification of Etfαβ sequences (Garcia Costas et al., 2017). Multiple protein sequence alignments were conducted using MUSCLE version 5.1, utilizing its default algorithm. The ML phylogenetic tree was generated employing the Le_Gascuel_2008 model, incorporating a discrete Gamma distribution to account for evolutionary rate differences among sites. Bootstrap values were calculated from 500 replicates to assess the robustness of the tree. Comprehensive details on all genome accession numbers, the phylogeny of the analyzed genomes, and locus tags for Etfα- and β-like genes are provided in **Dataset S2**.

**Quantification of *apmAB* in the fecal samples through quantitative PCR (qPCR)**

The copy number of *apmAB* in the fecal DNA samples was determined using qPCR methods as described in a previous study (Hsiao et al., 2023). The calibration curves were obtained by using 10-fold serial dilution of the full-length PCR product of *C. innocuum apmA* and *apmB*, respectively (**Figure S10**). The primer sets used for full-length *apmAB* cloning and qPCR were listed in the Table S5.

**Determination of the expression of strain RGG8 genes under different growth conditions**

The RNA was extracted from RGG8 cells treated with different chemicals (e.g., quinoline, menadione, progesterone, and glucose) using Direct-zol™ RNA Miniprep kit (Zymo Research, Irvine, CA, USA); the complementary DNA was generated using SuperScript™ IV First-Strand Synthesis System (Thermo Fisher Scientific, Waltham, MA, USA) as described in a previous study (Hsiao et al., 2023). The relatively expressional level of *apmA* in the different treatments was evaluated by using the $2^{-\Delta\Delta Ct}$ method with the Ct value of bacterial universal 16S rRNA (see **Table S5** for nucleotide sequences) as the internal control. The expression of *apmA* in the control groups (DMSO or 0.05% glucose) was set as 1.

**PacBio sequencing of bacterial 16S rRNA gene amplicons**

Genomic DNA of bacteria was extracted using a column-based kit (QIAamp PowerFecal DNA Kit, Qiagen). DNA concentration was determined using a Qubit 4.0 Fluorometer (Thermo Scientific). The full-length of bacterial 16S rRNA gene (V1-V9 regions) was amplified using the barcoded 16S gene-specific universal primer set. According to the PacBio standard protocol (Amplification of full-length 16S gene with barcoded primers for multiplexed SMRTbell library preparation and sequencing procedure), each primer is designed to include a 5' buffer sequence (GCATC) with a 5' phosphate modification, a 16-base barcode, and the degenerate 16S gene-specific universal forward or reverse primer sequences. (Forward:5'Phos/GCATC- 16-base barcode -AGRGTTYGATYMTGGCTCAG -3', Reverse: 5'Phos/GCATC- 16-base barcode – RGYTACCTTGTTACGACTT -3'). Degenerate base identities are M = A, C; R = A, G; and Y = C, T. In brief, 2 ng of gDNA was used for the PCR reaction carried out with KAPA HiFi HotStart ReadyMix (Roche) under the PCR condition: initial denaturation at 95°C for 3 min; followed by 20 to 30 cycles (sample dependence) of: 95°C for 30 sec, 57°C for 30 sec, 72°C for 60 sec; a final elongation step for 5 min at 72°C and then incubated at 4°C. The success of each PCR amplification was examined on 1% agarose gel. Samples with a bright band around 1,500bp were chosen and purified using the AMPure PB Beads for the downstream PacBio library preparation.

**Bacterial community structure analysis**

The raw PacBio reads were quality-filtered and analyzed using QIIME2-DADA2 pipeline (ver. 2023.5) (Hall & Beiko, 2018). Low-quality sequences were removed from the PacBio raw reads (i.e., sequences with a length lower than 1000 bp or longer than 1600 bp, primer and chimeric sequences), dereplicated, and amplicon sequence variants (ASVs) with a near-zero error rate and single-nucleotide resolution were generated. The ASVs were classified to the species level based on bacteria 16S rRNA RefSeq sequences from the NCBI nucleotide database (https://ftp.ncbi.nlm.nih.gov/refseq/TargetedLoci/Bacteria/). ASVs with sequence similarity higher than 98% were classified to the species level. Each sample was rarefied (i.e., normalized) to an equal sequencing depth (i.e., 12674 reads). The number of each ASV in each sample was transformed into relative abundance (i.e., proportion; %).

**Preparation of strain RGG8 cell suspension for mice administration**

Strain RGG8 was anaerobically grown in Gifu Anaerobic Broth (600 mL), and the bacterial cultures were incubated at 37°C in an orbital shaker (120 rpm) for approximately 16 hours. The bacterial cells were harvested through centrifugation, and the cell pellet was resuspended in an anaerobic phosphate buffer saline (15 mL). The colony-formation-units (CFUs) of the resulting cell suspension were determined by counting the numbers of strain RGG8 colonies grown on Gifu Anaerobic agar. This cell suspensions of strain RGG8 were stored at 4 °C (within 5 days) before use.

**Strain RGG8 administration through oral gavage**

C57BL/6J mice (aged 7-week-old) were obtained from the Animal Center of the Medical College (National Taiwan University, Taipei, Taiwan) and were kept in standard animal housing conditions with the health guidelines for the care and use of experimental animals. The experiments were approved by the local ethics committee (IACUC No. 20220423). After acclimatization (including estrous cycle synchronization by using male mice urine and microbiota removal by using administration of ABX solution [Amphotericin-B 0.1mg/ml; Ampicillin 10mg/ml; Neomycin 10mg/ml; Metronidazole 10mg/ml; Vancomycin 5mg/ml] through oral-gavage) for one week, mice with similar body weight (16–18 g) were randomly assigned into the treatment groups as the descriptions in the main text (**Figures 3Ai and 3Bi**). For administration of stain RGG8, approximately 5 × $10^8$ CFUs (suspended in 200 uL of basal mineral medium) were fed into each mouse through oral gavage twice per week. For exogenous progesterone administration by oral gavage, mice were orally administered with Utrogestan (OLIC (THAILAND) LIMITED) (20 mg/kg/day) suspended in sesame oil (Sigma-Aldrich, St. Louis, MO, USA). Fresh mouse feces were collected daily, and the body weight was measured once per week. Mice were sacrificed 4 weeks or 12 weeks (anesthetized with 3% isoflurane) and the mouse blood was collected through cardiac puncture. The ovaries were excised, weighed, fixed in neutral-buffered 4% formaldehyde for 24 hours, washed with distilled water, dehydrated and embedded in paraffin. Serum samples were stored at -80°C before use.

**Vaginal smears for estrus phase determination**

The stage of the estrous cycle was determined by daily vaginal smears taken 10 series day before the mice were euthanized. The vaginal smear was performed according to the following procedure as described in a previous study (Chen et al., 2016). In brief, a small amount of a saline solution was inserted into the mouse vagina with a disposable pipette, removed, placed on a slide and examined under a microscope. The average interestrus interval was defined as the average duration of the dominant presence of characteristic non-nucleated, cornified epithelial cells with a high cell density during the 10 days of experiments for vaginal smears.

**Ovarian histology and antral follicle counts**

The ovaries were serially sectioned in the long axis at 5-μm intervals and every fifth section was stained with hematoxylin and eosin (HE). The section with the largest diameter of each ovary was chosen as the representative section for that ovary. For each representative section, counts of the total number of follicles and number of follicles at each developmental stage were determined as described in a previous study (Yang et al., 2021). All the follicles with intact, non-fragmented oocytes on the representative sections were counted and classified by developmental stage. The developmental stage for a particular follicle was determined after reviewing all the adjacent sections containing the follicle. All the histopathology counts and classifications of the follicles were performed by two observers who reviewed the slides in collaboration before finalizing the counts. Both observers were blinded to the treatment group. The secondary follicle was characterized by two or more layers of granulosa cells without a visible antrum, and a tertiary follicle was identified by the presence of an antrum. Tertiary follicles were further divided into early and late tertiary follicles based on the projected oocyte maturity. Pre-ovulatory (Graafian) follicles and follicles with diameters ≥ 250 μm, a criterion reported to be a good indicator of oocyte maturity in mice were categorized as late tertiary follicles. Tertiary follicles without these developmental features were categorized as early tertiary follicles. The secondary follicles were counted at 400x magnifications, and the tertiary follicles were counted at 40x magnifications.

**Determination of serum progesterone levels**

Serum progesterone levels of enrolled women with infertility were measured using indirect chemiluminescence (VitrosEci; Ortho Clinical Diagnostics, Rochester, NY) (Yang et al., 2018). The serum progesterone level of mice was determined using Crystal Chem's Mouse Progesterone ELISA Kit (Catalog# 80559, Crystal Chem, IL, USA) and was performed following the manufacturer's instructions.

**Participants and protocols of hormone treatment and samples collection**

A total of 14 infertile women undergoing hormone treatment for endometrial preparation for frozen-thaw embryo transfer were included in this study. The endometrium was primed through the administration of exogenous estrogen and progesterone. Hormone replacement therapy commenced on day 3 of the menstrual cycle, with estradiol valerate administered at 4 mg/day from day 3 to 8, 8 mg/day from day 9 to 11, and 12 mg/day from day 12 onward until achieving the optimal endometrial thickness for embryo transfer. On the day when the embryo transfer was scheduled, 8% progesterone gel (Crinone®, Merck) at a dosage of 90 mg/day was administered transvaginally for 2 days, followed by an increased dosage of 180 mg/day for the subsequent 14 days. Oral progesterone (Utrogestan) at a dosage of 600 mg/day was initiated on the day of the embryo transfer. All participants were required to provide fecal samples on the day preceding the commencement of progesterone treatment. Serum samples, obtained three to four times, were collected on the day prior to initiating progesterone treatment, two days after commencing oral progesterone, and one week later. In case of a positive pregnancy test result, an additional blood sample would be collected one week after the confirmation of pregnancy. The procedure was approved by the local ethics committee (Clinical Trial /Research Approval NTUH-REC No.:202103046RINB).

### Supplemental Figures

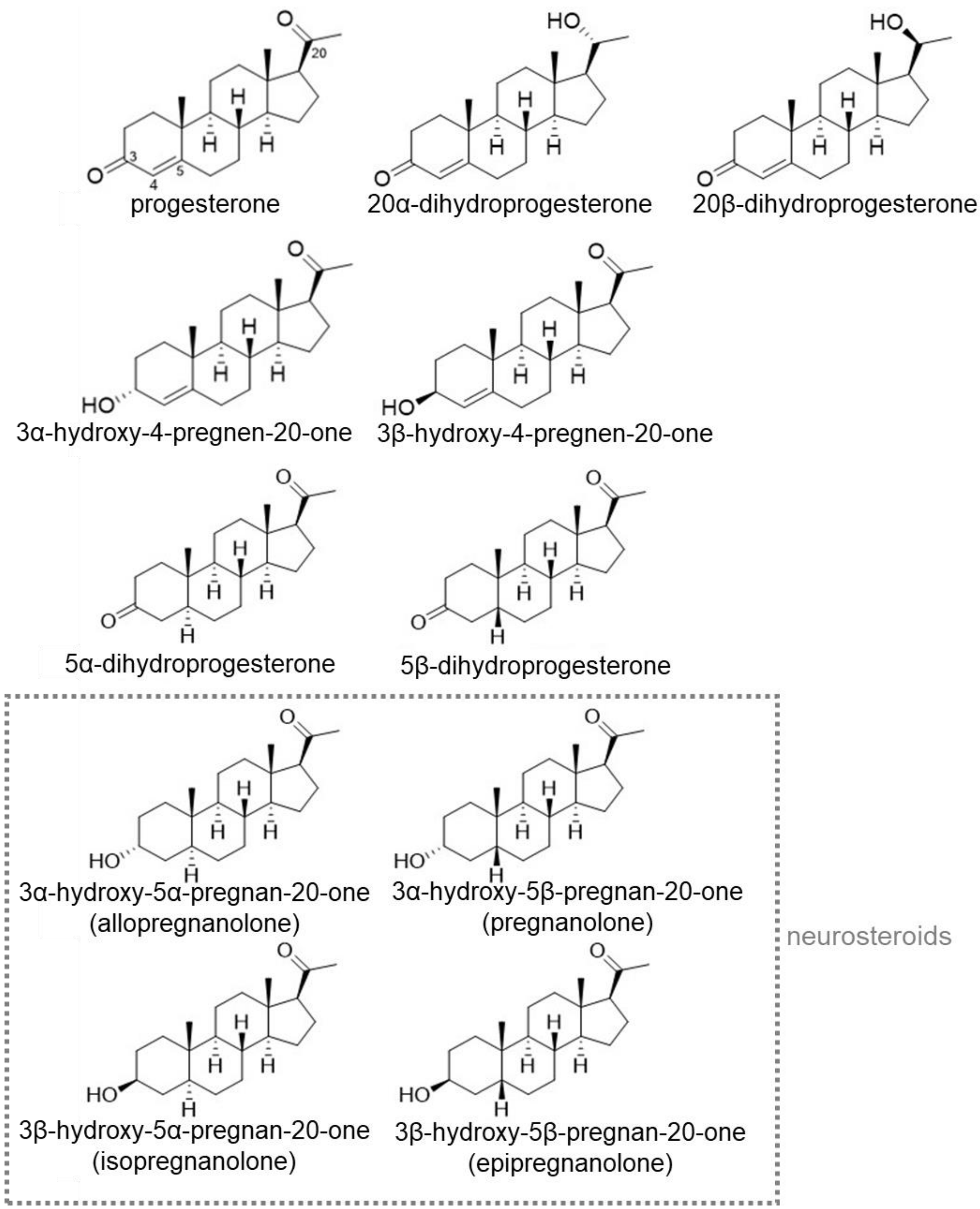


**Fig. S1.** The chemical structures of prevalent progestogens. Among them, pregnanolone, allopregnanolone, epipregnanolone, and isopregnanolone are considered neurosteroids. Critical carbons in the steroidal numbering system are shown on progesterone.

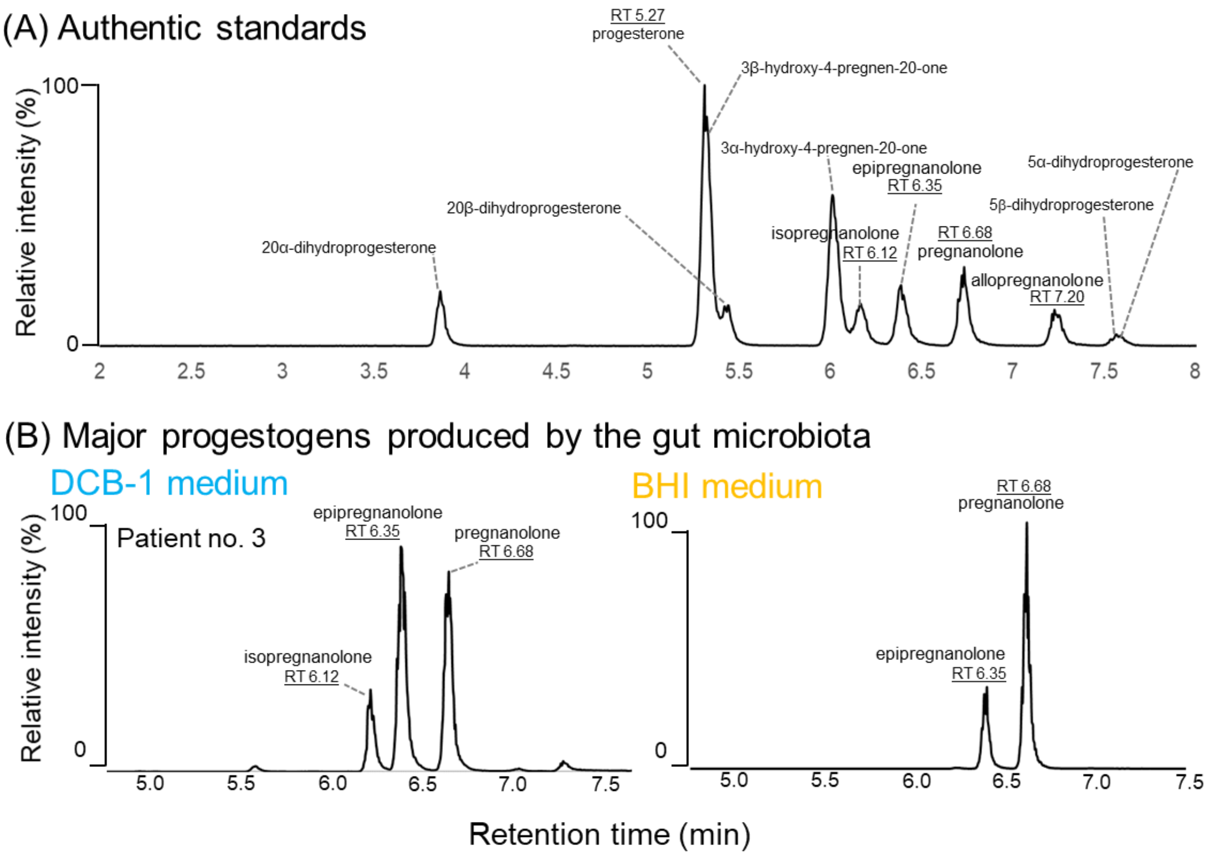


**Fig. S2.** Identification of major progestogenic metabolites produced by the gut microbiota of Patient no. 1. (A) The UPLC chromatogram of individual progestogenic standards. Abbreviation: 3β5α, 3β-hydroxy-5α-pregnan-20-one. (B) Major progestogenic metabolites produced by the gut microbiota of Patient no. 1. Both pregnanolone and epipregnanolone were produced in the fecal cultures anaerobically incubated with progesterone (1 mM) in the DCB-1 mineral medium or the BHI rich medium.

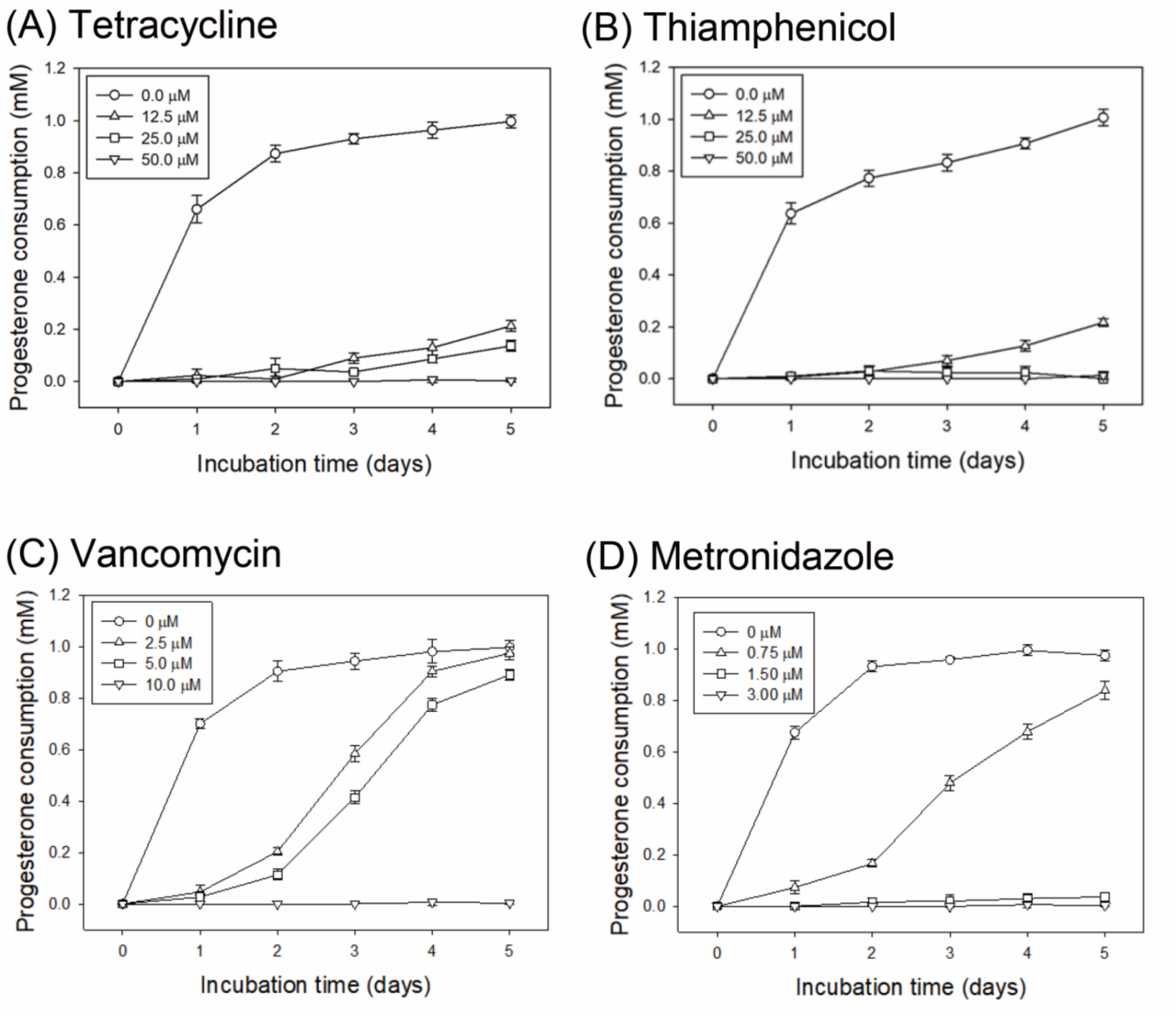


**Fig. S3.** Effect of two broad-spectrum antibiotics tetracycline (**A**) and thiamphenicol (**B**) and two *Clostridium*-specific antibiotics vancomycin (**C**) and metronidazole (**D**) on the anaerobic progesterone metabolism by the gut microbiota from Patient no. 1.

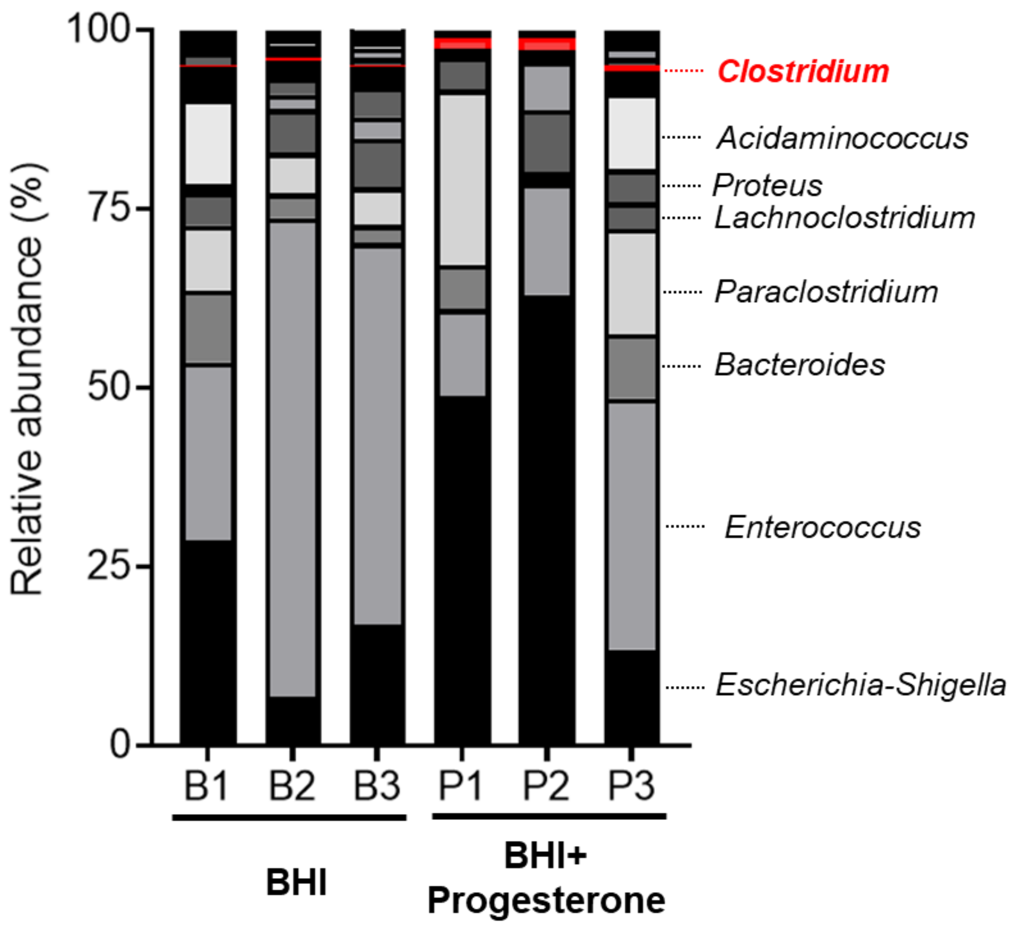


**Fig. S4.** Impact of progesterone administration on the gut microbiota**.** Gut bacterial communities across the different treatments (BHI broth incubated with or without 1 mM progesterone; triplicates) was analyzed by sequencing the bacterial 16 rRNA amplicons on a PacBio platform.

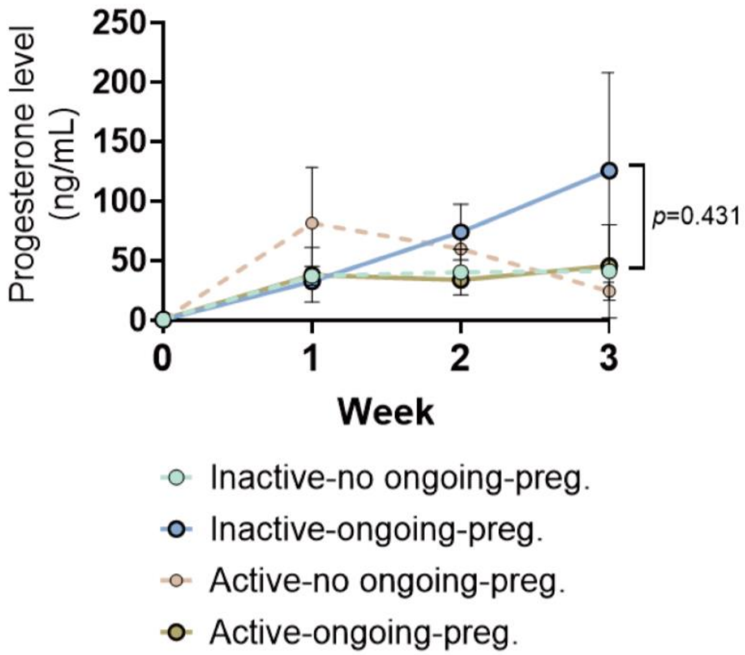


**Fig. S5.** Trend of serum progesterone level of the pregnant patients who received the embryo transfer. The criterion of subgrouping the patients into ongoing or no ongoing pregnancy was determined by the observation of gestational sac after the embryo transfer for 3 weeks. The data was presented as the mean ± SE.

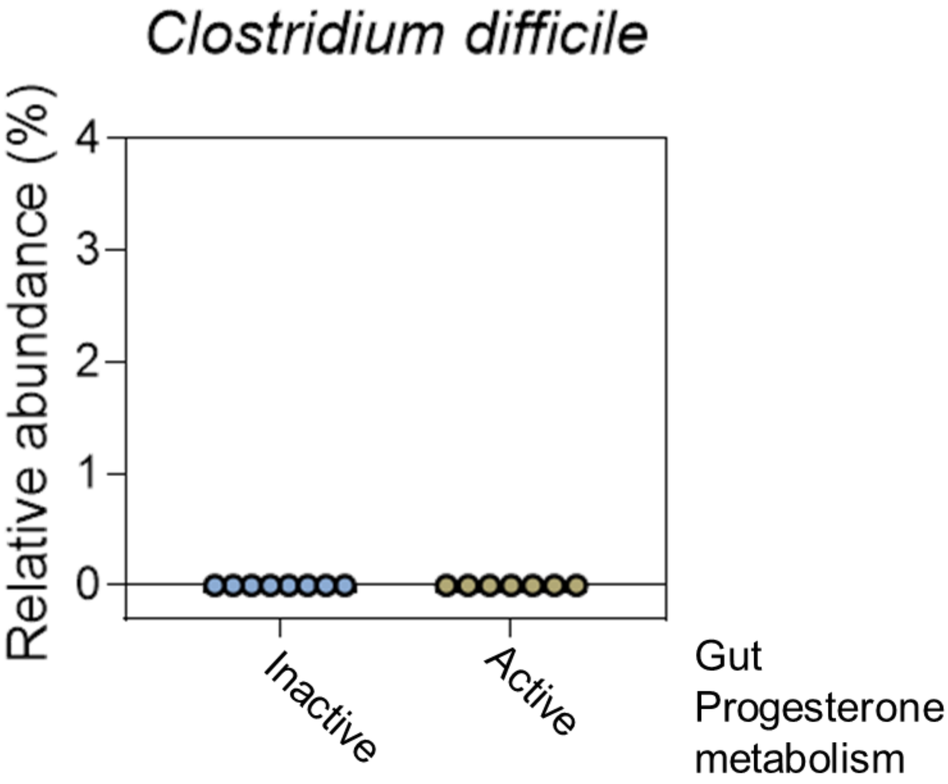


**Fig. S6.** Relative abundance of *Clostridium difficile* in the active gut microbiota (capable of progesterone metabolism). Totally 14 gut microbiota from infertile women were tested. Among them, eight gut microbiota exhibited negligible progesterone metabolic activity (Inactive group; n = 8), whereas the remaining gut microbiota exhibited apparent progesterone metabolic activity (Active group; n = 6).

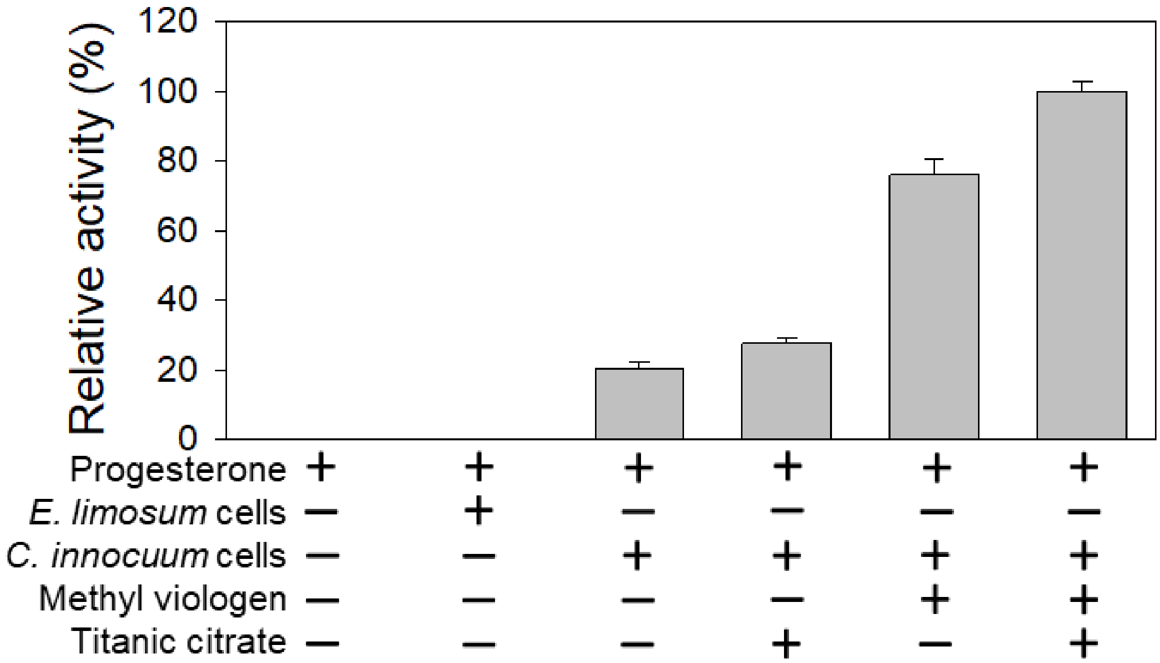


**Fig. S7.** Resting cell assays indicated the extracellular electron carrier requirement and oxygen sensitivity for the progesterone metabolism by *C. innocuum* strain RGG8 cells.

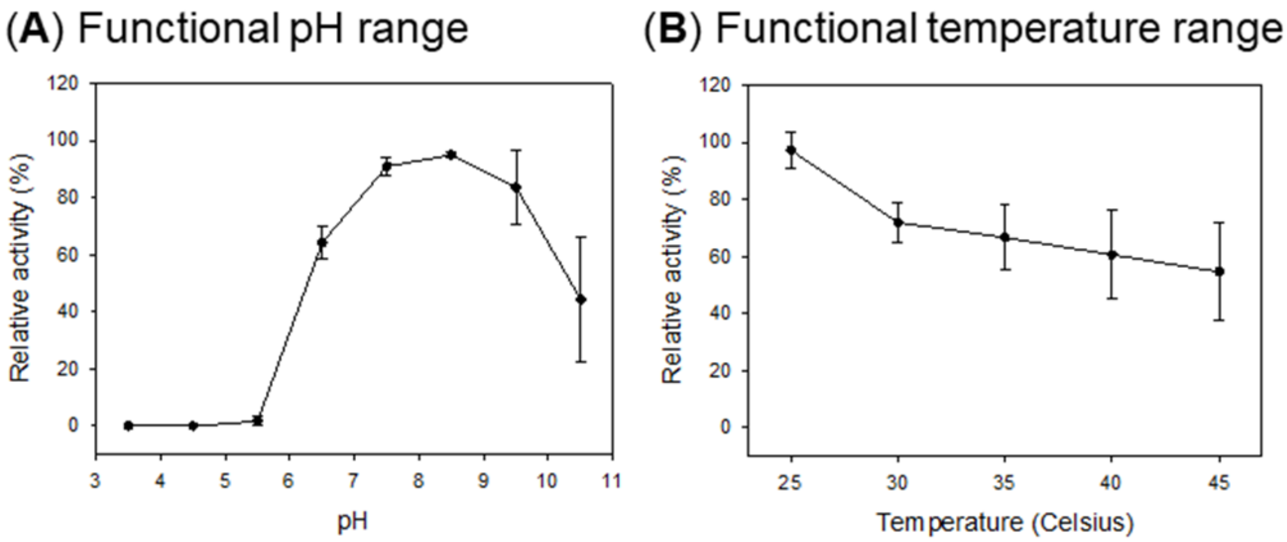


**Fig. S8.** Opitmal working pH and temperature of the 5β-dihydroprogesterone reductase purified from *C. innocuum* strain RGG8.

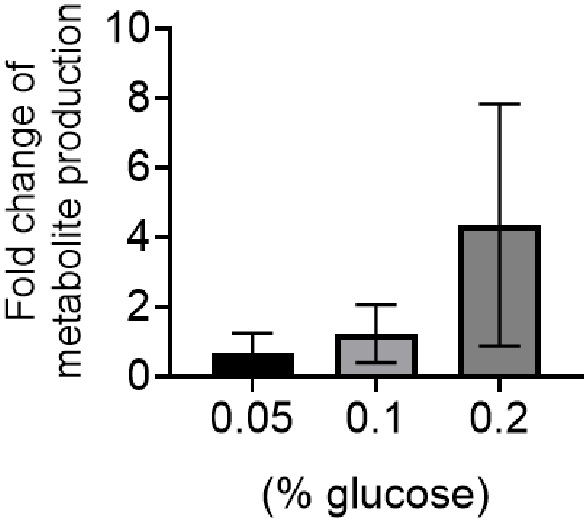

**Fig. S9.** Glucose induced the production of epipregnanolone by *Clostridium innocuum* strain RGG8. The metabolite production was examined by using thin-layer chromatography from 3 individual experiments. The data was presented by mean ± SE.

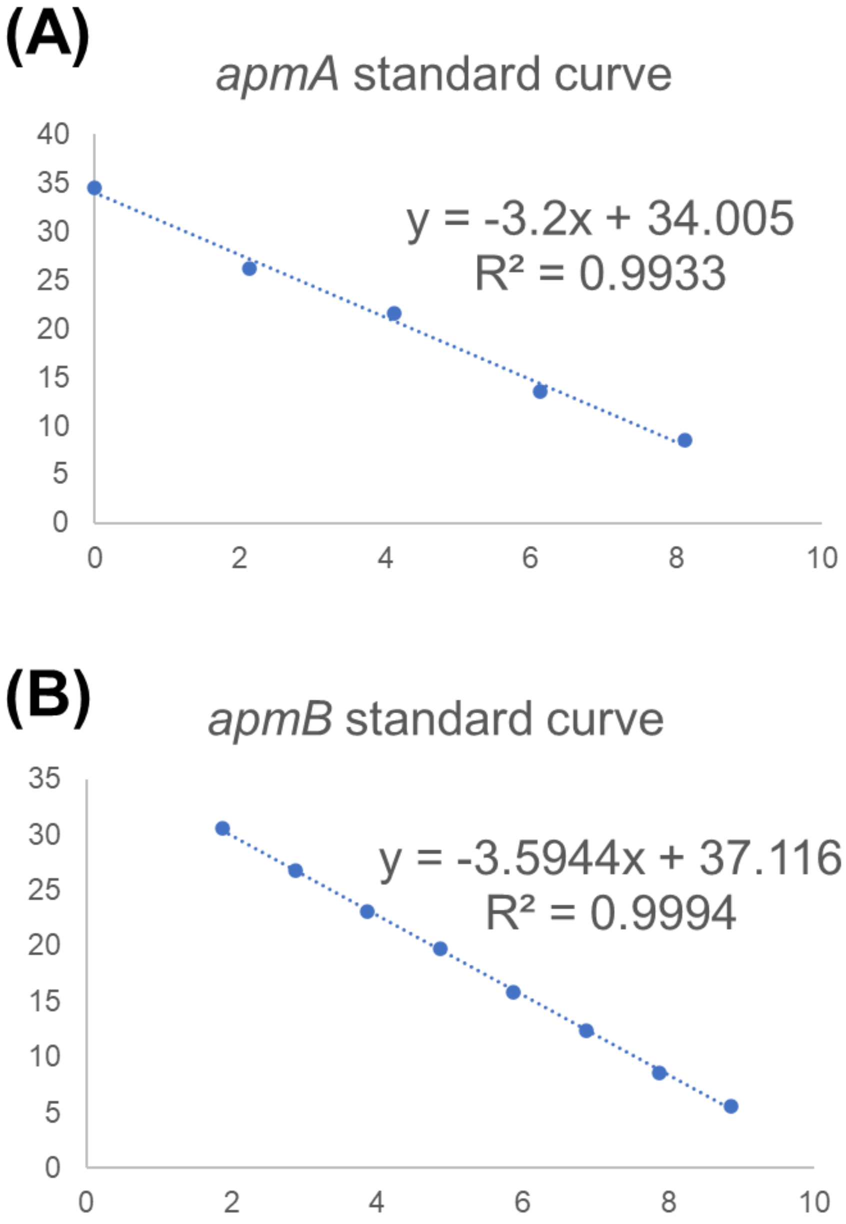


**Fig. S10.** The qPCR standard curves of primer pairs specific for *apmA* ($R^2$ = 0.99) and *apmB* ($R^2$ = 1.00).

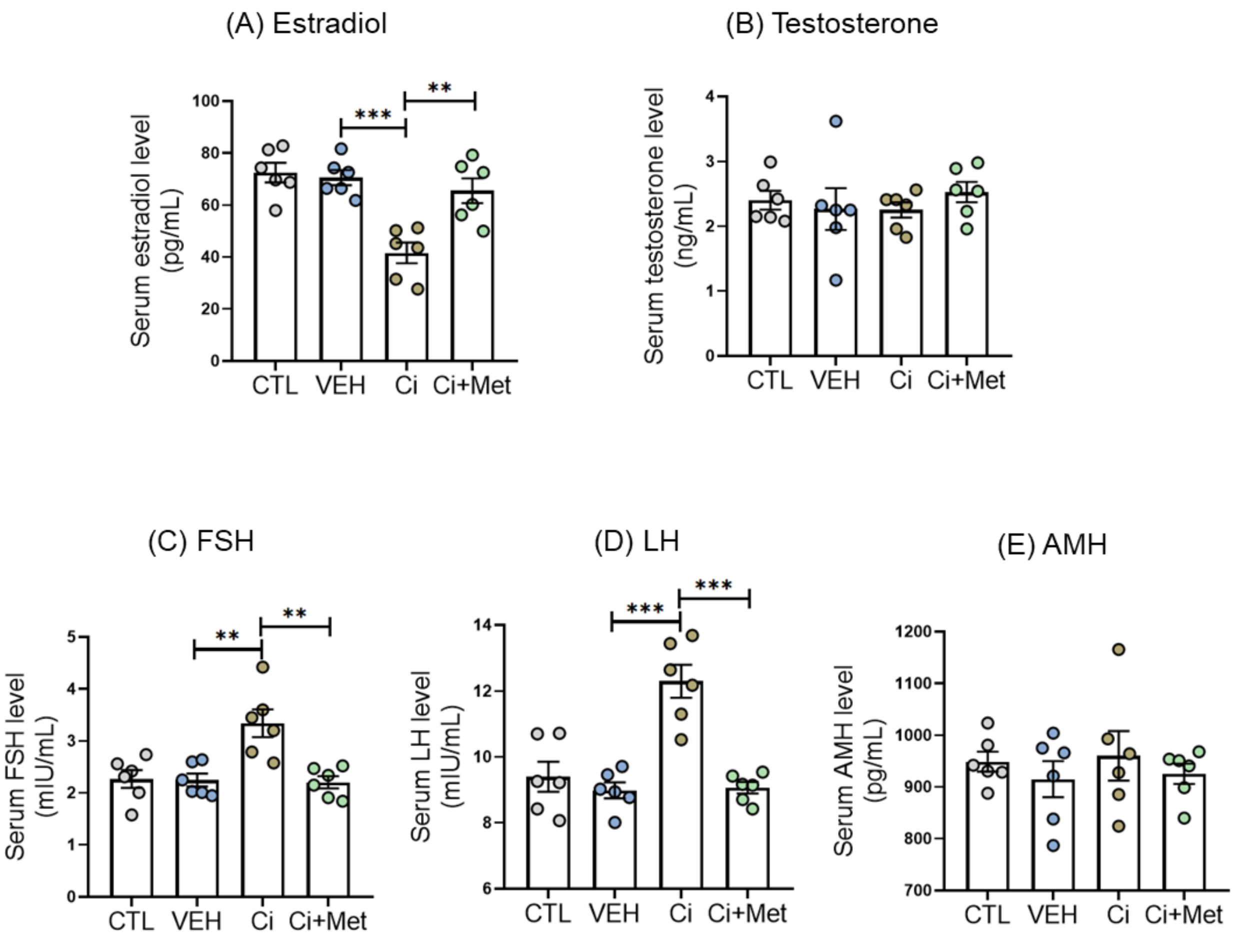


**Fig. S11.** The *C. innocuum* administration cause a significant decrease in mouse serum estradiol level (A) but did not change in serum testosterone level (B). Moreover, the *C. innocuum* administration appeared to elevate the serum FSH (C) and LH levels (D) but did not lead to apparent changes in the serum AMH level (E).

### Supplemental Tables

**Table S1.** Physiological characteristics of 14 infertile patients.

| Patient No | Progesterone metabolic activity* | Age (years) | BMI ($Kg/m^2$) | Indication for IVF | Menstrual regularity | AMH level (ng/ml) | Pregnancy outcome |
|---|---|---|---|---|---|---|---|
| **Case 1** | active | 36.2 | 36.7 | Primary infertility and tubal factor | irregular | 1.76 | live birth |
| **Case 2** | inactive | 35.9 | 30.9 | Primary infertility | irregular | 6.52 | chemical pregnancy |
| **Case 3** | inactive | 36.8 | 27.2 | Secondary infertility | regular | NA | chemical pregnancy |
| **Case 4** | inactive | 33.3 | 24.3 | Primary infertility and male factor | regular | 4.82 | live birth |
| **Case 5** | inactive | 35.4 | 20.5 | Primary infertility | regular | 4.03 | no pregnancy |
| **Case 6** | inactive | 35.9 | 21.7 | Primary infertility and male factor | regular | 1.31 | no pregnancy |
| **Case 7** | active | 38.9 | 31.3 | Primary infertility | regular | 0.62 | chemical pregnancy |
| **Case 8** | inactive | 34.5 | 21.1 | Primary infertility and premature ovarian failure | irregular | <0.02 | live birth |
| **Case 9** | inactive | 37.4 | 21.6 | Primary infertility and tubal factor | regular | 3.78 | no pregnancy |
| **Case 10** | active | 37.4 | 19.2 | Secondary infertility | irregular | NA | chemical pregnancy |
| **Case 11** | active | 32.2 | 20.3 | Secondary infertility and tubal factor | regular | 2.65 | chemical pregnancy |
| **Case 12** | inactive | 42.6 | 22.5 | Primary infertility and tubal factor | regular | 2.57 | live birth |
| **Case 13** | active | 40.4 | 23.8 | Secondary infertility and male factor | regular | 4.38 | live birth |
| **Case 14** | active | 40.4 | 39.5 | Primary infertility and tubal factor | irregular | 6.38 | live birth |

NA: not available

*Progesterone metabolic activity was determined by quantifying residual progesterone remaining in fecal cultures that were anaerobically incubated with progesterone (1 mM) in the DCB-1 medium for 7 days. “Active” is represented by the microbial utilization of > 50% of progesterone molecules.

**Table S2.** UPLC-HRMS behaviors of individual progestogenic standards.

| Compound | Chemical formula | Molecular Weight | Adduct *m/z* | RT (min) |
|---|---|---|---|---|
| 5β-dihydroprogesterone | $C_{21}H_{32}O_2$ | 316.48 | 317.2469 $[M+H]^+$ | 7.56 |
| 5α-dihydroprogesterone | $C_{21}H_{32}O_2$ | 316.48 | 317.2472 $[M+H]^+$ | 7.59 |
| 3α-hydroxy-4-pregnen-20-one | $C_{21}H_{32}O_2$ | 316.48 | 299.2362 $[M-H_2O+H]^+$ | 6.02 |
| 3β-hydroxy-4-pregnen-20-one | $C_{21}H_{32}O_2$ | 316.48 | 299.2368 $[M-H_2O+H]^+$ | 5.31 |
| 20α-dihydroprogesterone | $C_{21}H_{32}O_2$ | 316.48 | 317.2468 $[M+H]^+$ | 3.86 |
| 20β-dihydroprogesterone | $C_{21}H_{32}O_2$ | 316.48 | 317.2471 $[M+H]^+$ | 5.43 |
| 3β-hydroxy-5α-pregnan-20-one (isopregnanolone) | $C_{21}H_{34}O_2$ | 318.49 | 301.2523 $[M-H_2O+H]^+$ | 6.12 |
| 3β-hydroxy-5β-pregnan-20-one (epipregnanolone) | $C_{21}H_{34}O_2$ | 318.49 | 301.2522 $[M-H_2O+H]^+$ | 6.35 |
| 3α-hydroxy-5α-pregnan-20-one (allopregnanolone) | $C_{21}H_{34}O_2$ | 318.49 | 301.2522 $[M-H_2O+H]^+$ | 7.20 |
| 3α-hydroxy-5β-pregnan-20-one (pregnanolone) | $C_{21}H_{34}O_2$ | 318.49 | 301.2522 $[M-H_2O+H]^+$ | 6.68 |
| Progesterone | $C_{21}H_{30}O_2$ | 314.22 | 315.2423 $[M+H]^+$ | 5.27 |

**Table S3.** Determination of the abundance of the progesterone metabolic genes *apmAB* in individual fecal samples through qPCR.

| Patient no. | Progesterone metabolic activity* | *apmA* abundance | *apmB* abundance |
|---|---|---|---|
| Case 1# | active | $1.56 \times 10^7$ | $2.56 \times 10^7$ |
| Case 2 | inactive | $0.47 \times 10^7$ | $0.9 \times 10^7$ |
| Case 3 | inactive | $0.002 \times 10^7$ | 0 |
| Case 4 | inactive | 0 | 0 |
| Case 5 | inactive | $0.003 \times 10^7$ | 0 |
| Case 6 | inactive | $0.04 \times 10^7$ | $0.04 \times 10^7$ |
| Case 7 | active | 0 | 0 |
| Case 8 | inactive | $0.01 \times 10^7$ | 0 |
| Case 9 | inactive | $0.12 \times 10^7$ | $0.17 \times 10^7$ |
| Case 10 | active | $0.13 \times 10^7$ | $0.24\times 10^7$ |
| Case 11 | active | $4.05 \times 10^7$ | $5.8 \times 10^7$ |
| Case 12 | inactive | $0.08 \times 10^7$ | $0.23 \times 10^7$ |
| Case 13 | active | $4.35 \times 10^7$ | $6.13 \times 10^7$ |
| Case 14 | active | $0.05 \times 10^7$ | $0.099 \times 10^7$ |

#, *C. innocuum* strain RGG8 was isolated from the gut microbiota of Patient no. 1.

*, Progesterone metabolic activity was determined by quantifying residual progesterone remaining in fecal cultures that were anaerobically incubated with progesterone (1 mM) in the DCB-1 medium for 7 days. "Active" is represented by the microbial utilization of > 50% of progesterone molecules.

**Table S4.** The isolation and characterization of *Clostridium* species from the fecal samples of the Patient no. 1.

| Bacterial Identities | | Progesterone transformation | | progesterone metabolic genes | | Antibiotics susceptibility | | |
|---|---|---|---|---|---|---|---|---|
| *Clostridium* species | Strains | Epipregnanolone production | Pregnanolone production | *apmA* (LMAICMKE_01819) | *apmB* (LMAICMKE_01820) | Metronidazole (broth) (MIC; μg/ mL ) | Metronidazole (agar) (MIC; μg/ mL) | Vancomycin (broth) (MIC; μg/ mL ) |
| *C. innocuum* | RGG8 | +++ | + | + | + | 1.25 | 0.30 | R |
| | RBW1-1 | ++ | + | + | + | 1.25 | 0.30 | R |
| | BT6 | ++ | + | + | + | 1.25 | 0.30 | R |
| | BP16 | ++ | + | + | + | 1.25 | 0.30 | R |
| | ATCC14501 | ++ | + | + | + | 1.25 | 0.30 | R |
| *C. difficile** | BCRC17678 | - | - | - | - | 1.25 | 0.25 | 2.50 |
| | BCRC17702 | - | - | - | - | 1.25 | 0.25 | 2.50 |
| | BCRC17900 | - | - | - | - | 1.25 | 0.25 | 2.50 |
| | BCRC80997 | - | - | - | - | 1.25 | 0.25 | 2.50 |
| | BCRC80998 | - | - | - | - | 1.25 | 0.25 | 1.25 |
| | BCRC80999 | - | - | - | - | 0.63 | 0.25 | 1.25 |
| *C. butyricum* | M3 | - | - | - | - | 0.31 | 0.10 | 1.25 |
| | C14-1 | - | - | - | - | 0.31 | 0.10 | 1.25 |
| | C14-4 | - | - | - | - | 0.31 | 0.10 | 1.25 |
| | C14-50 | - | - | - | - | 0.31 | 0.10 | 1.25 |
| | C14-51 | - | - | - | - | 0.31 | 0.10 | 1.25 |
| *C. scindens* | L1-10 | - | - | - | - | 0.15 | 0.70 | 2.50 |
| | L1-28 | - | - | - | - | 0.15 | 0.70 | 2.50 |
| | L1-33 | - | - | - | - | 0.15 | 0.70 | 2.50 |
| *C. subterminale* | L2B4-2a | - | - | - | - | R | R | 1.25 |
| *C. tertium* | L2D7-4 | - | - | - | - | R | R | 1.25 |
| *C. acetobutylicum* | L3D7-7 | - | - | - | - | 0.15 | 0.10 | 1.25 |

*, strains of *C. difficile* was obtained from Bioresource Collection and Research Center, Food Industry Research and Development Institute, Hsinchu, Taiwan as the gut microbiota of Patient no. 1 did not contain *C. difficile*.
+++, > 80% of the substrate was transformed into corresponding product; ++, > 50% (but < 80%) of the substrate was transformed into corresponding product; +, > 10% (but < 50%) of the substrate was transformed into corresponding products.

**Table S5.** Nucleotide sequences of the PCR primers used in this study.

| Primer | Sequence (5′- 3′) | Usage | **Reference** |
|---|---|---|---|
| *apmA* full-length primer F | ATGAAAATTATCGTTCTTGTAAAAC | Amplifying *apmA* gene (full-length) from strain RGG8 DNA | This study |
| *apmA* full-length primer R | CTATTTCGTAATAATCTGCTTGTC | Amplifying *apmA* gene (full-length) from strain RGG8 DNA | This study |
| *apmA* qPCR primer F | ACATTGATTCGTGCCGGAGT | Quantifying the expression of strain RGG8-specific *apmA* | This study |
| *apmA* qPCR primer R | TTCAGCATTCCCGTAGCCTG | Quantifying the expression of strain RGG8-specific *apmA* | This study |
| *apmB* full-length primer F | ATGGCTAAATTTGAAGGATATAAA | Amplifying *apmB* gene (full-length) from strain RGG8 DNA | This study |
| *apmB* full-length primer R | TTATCCCTTTTTCGCAGC | Amplifying *apmB* gene (full-length) from strain RGG8 DNA | This study |
| *apmB* qPCR primer F | CACGTTTGAATGCCGGTCTG | Quantifying the expression of strain RGG8-specific apmB | This study |
| *apmB* qPCR primer R | CCATCTGCGGACGAGTATCC | Quantifying the expression of strain RGG8-specific apmB | This study |
| Bacteria 16s universal primer 27F | AGAGTTTGATCCTGGCTCAG | Amplifying bacteria 16s rRNA gene from total DNA extracts | Miller et al. (2013) |
| Bacteria 16s universal primer 1492R | GGTTACCTTGTTACGACTT | Amplifying bacteria 16s rRNA gene from total DNA extracts | Miller et al. (2013) |

### Legends for Datasets

**Dataset S1 (separate file).** Genome annotation of the strain RGG8.

**Dataset S2 (separate file).** Genes of the Etf protein family selected for phylogenetic analysis.
